# Loop-extruding Smc5/6 organizes transcription-induced positive DNA supercoils

**DOI:** 10.1101/2023.06.20.545053

**Authors:** Kristian Jeppsson, Biswajit Pradhan, Takashi Sutani, Toyonori Sakata, Miki Umeda Igarashi, Davide Giorgio Berta, Takaharu Kanno, Ryuichiro Nakato, Katsuhiko Shirahige, Eugene Kim, Camilla Björkegren

## Abstract

The Structural Maintenance of Chromosome (SMC) protein complexes cohesin, condensin and the Smc5/6 complex (Smc5/6) are essential for chromosome function. At the molecular level, these complexes fold DNA by loop extrusion. Accordingly, cohesin creates chromosome loops in interphase, and condensin compacts mitotic chromosomes. However, the role of Smc5/6’s recently discovered DNA loop extrusion activity is unknown. Here, we uncover that Smc5/6 associates with transcription-induced positively supercoiled chromosomal DNA at cohesin-dependent chromosome loop boundaries. Mechanistically, single-molecule imaging reveals that dimers of Smc5/6 specifically recognize the tip of positively supercoiled DNA plectonemes, and efficiently initiates loop extrusion to gather the supercoiled DNA into a large plectonemic loop. Finally, Hi-C analysis shows that Smc5/6 links chromosomal regions containing transcription-induced positive supercoiling in *cis*. Altogether, our findings indicate that Smc5/6 controls the three-dimensional organization of chromosomes by recognizing and initiating loop extrusion on positively supercoiled DNA.

## INTRODUCTION

The Smc5/6 complex (Smc5/6) is part of the eukaryotic family of Structural Maintenance of Chromosomes (SMC) protein complexes, which also includes cohesin and condensin. These multi-subunit complexes hydrolyze ATP to create DNA loops by extrusion (Davidson *et al*., 2019; Ganji *et al*., 2018; Pradhan *et al*., 2023). Cohesin performs so called symmetrical extrusion, *i.e.*, reels in DNA from both sides of a growing loop, while condensin creates loops by one-sided extrusion (Davidson *et al*., 2019; Ganji *et al*., 2018). Similar to cohesin, Smc5/6 performs two-sided loop extrusion. However, unlike cohesin and condensin, which extrude loops as monomers, Smc5/6 extrudes loops in the form of dimers of complexes (Pradhan *et al*., 2023). In line with loop extrusion taking place *in vivo*, cohesin promotes chromosome loop formation and formation of topological associated domains (TADs), and condensin compacts mitotic chromosomes (Gibcus *et al*., 2018; Rao *et al*., 2017; Schwarzer *et al*., 2017; Wutz *et al*., 2017). It remains, however, unknown if Smc5/6 loop extrusion activity also regulates the three-dimensional structure of chromosomes.

Smc5/6 cellular function remains mostly unknown, but appears to be executed during late S to G2/M-phase and prevents the formation of segregation-inhibiting chromatid linkages (Menolfi *et al*., 2015; Torres-Rosell *et al*., 2007; Venegas *et al*., 2020). Smc5/6 is also needed for DNA damage repair and controls homologous recombination (Irmisch *et al*., 2009; Lindroos *et al*., 2006; Torres-Rosell *et al*., 2005). Smc5/6 co-localizes with cohesin in between convergently oriented gene pairs on budding yeast, *Saccharomyces cerevisiae* (*S. cerevisiae*) chromosomes (Jeppsson *et al*., 2014; Lengronne *et al*., 2004), which also limits the size of cohesin loops by creating boundaries that block the progression of the extruding complex (Costantino *et al*., 2020; Jeppsson *et al*., 2022). Interestingly, Smc5/6 chromosomal association depends on cohesin, whereas cohesin binding is not disrupted by inactivation of Smc5/6 (Jeppsson *et al*., 2014). The two complexes also co-localize between convergently oriented gene pairs (Jeppsson *et al*., 2014; Lengronne *et al*., 2004), which in turn limits the size of cohesin loops by creating boundaries that block the progression of the extruding complex (Costantino *et al*., 2020; Jeppsson *et al*., 2022). Both complexes also appear on chromosomes in early S-phase, show the most prominent binding in G2/M, and are removed at anaphase (Jeppsson *et al*., 2014; Lindroos *et al*., 2006; Michaelis et al., 1997). However, while cohesin is highly enriched between most convergently oriented gene pairs along chromosome arms, strong Smc5/6 association is only observed in centromere-proximal regions in wild-type cells (Jeppsson *et al*., 2014; Lindroos *et al*., 2006), indicating that cohesin influences Smc5/6 chromosomal association indirectly.

Smc5/6 function has also been connected to DNA supercoiling and sister chromatid intertwinings (SCIs). SCIs are formed if the replication machinery rotates with the turn of the DNA helix during replication. DNA supercoiling arises when translocating polymerases or helicases unwind the DNA double helix, for example during transcription (reviewed in (Wang, 2002)). This causes over-twisting of the DNA (*i.e.,* positive supercoiling) ahead, and under-twisting (*i.e.,* negative supercoiling) behind the transcription machinery (see Figure S1A-C for details). DNA supercoils and SCIs are resolved by type I and II topoisomerase enzymes that create transient single and double-strand DNA breaks, respectively. While SCIs are removed by type II topoisomerases (Top2), supercoil relaxation depends on both Top2 and type I topoisomerases (*e.g.*, Top1). When SCI resolution is inhibited by inactivation of Top2 during replication, Smc5/6 binding is no longer restricted to centromere-proximal regions but appear in between convergently oriented genes all along *S. cerevisiae* chromosome arms, thereby showing a more complete overlap with cohesin (Jeppsson *et al*., 2014; Kegel *et al*., 2011). This suggests that the presence of SCIs directly or indirectly controls the chromosomal association of Smc5/6. *In vitro* pull-down experiments show that Smc5/6 also associates to positively supercoiled plasmids, and magnetic tweezer analysis reveals that Smc5/6 stabilizes both positive and negative supercoils (Gutierrez-Escribano *et al*., 2020; Serrano *et al*., 2020). Altogether, this indicates that in addition to being regulated by cohesin, Smc5/6 function is connected to SCIs and supercoiling, but so far, the interrelationship between these three factors has remained undetermined.

Prompted by our recent finding that Smc5/6 is a DNA loop extruder *in vitro* (Pradhan *et al*., 2023), we have explored if Smc5/6 controls the spatial organization of chromosomes. This has revealed a specialized genome organization function where Smc5/6 creates intrachromosomal links between chromosomal regions that contain transcription-induced positive supercoiling. These regions are found at the base of cohesin-dependent loops, and loop-extruding cohesin is also found to control Smc5/6 chromosomal binding. Single-molecule imaging confirms the preferential binding of Smc5/6 to positive supercoils and further reveal that dimers of Smc5/6 initiate DNA loop extrusion at the tip of positively supercoiled DNA plectonemes. Altogether, our work establishes that all eukaryotic SMC complexes control the spatial organization of chromosomes, and reveals mechanistic insights into Smc5/6 cellular function and its connection to cohesin, SCIs, and DNA supercoiling.

## RESULTS

### Smc5/6 chromosomal association depends on loop-extruding cohesin

Given that Smc5/6 chromosomal localization depends on cohesin, and cohesin have been shown to create loops with boundaries at Smc5/6 binding sites, Smc5/6 might be controlled by loop-extruding cohesin (Costantino *et al*., 2020; Garcia-Luis *et al*., 2019; Jeppsson *et al*., 2014). To test this, we performed ChIP-seq and ChIP-qPCR analyses of Smc6 after depletion of Scc2 and Wpl1, the cohesin loader and unloader, respectively (Ciosk et al., 2000; Kueng et al., 2006; Tedeschi et al., 2013). When Scc2 is depleted in G2/M-arrested cells, the majority of cohesin-mediated loops are lost (Jeppsson *et al*., 2022), while sister chromatid cohesion, the canonical function of cohesin (Guacci *et al*., 1997; Michaelis *et al*., 1997), remains unperturbed (Ciosk *et al*., 2000). The analysis showed that Scc2 depletion in G2/M significantly reduces Smc5/6 enrichment around centromeres, indicating a dependency on cohesin loop extrusion (Figures 1A-C and S1D-E). Supporting this, Wpl1 depletion, which causes cohesin to remain on chromosomes and form more and longer loops, significantly increased Smc5/6 enrichment around centromeres (Figures 1A-C and S1D-E) (Jeppsson *et al*., 2022; Lopez-Serra *et al*., 2013).

**Figure 1.**
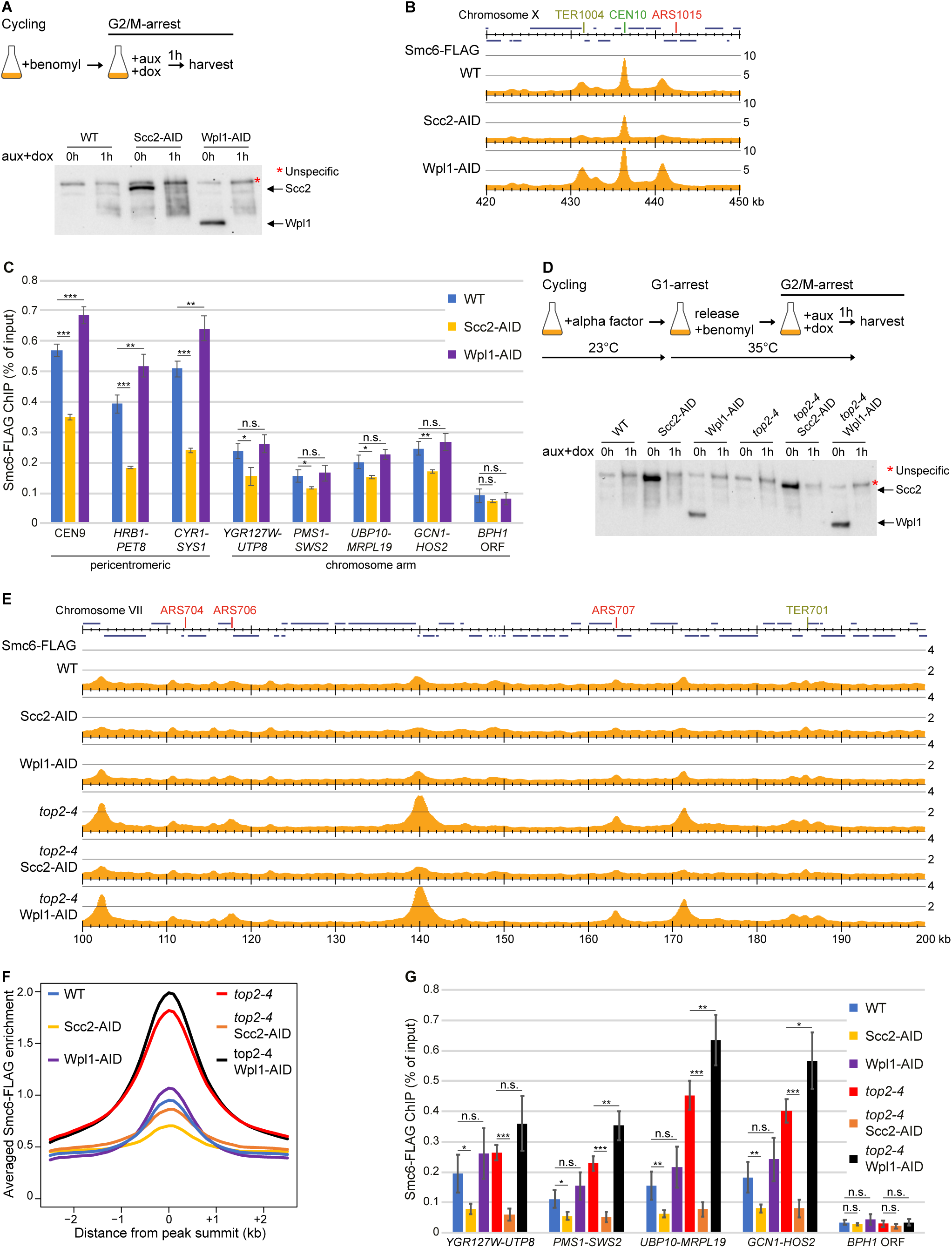
Smc5/6 chromosomal association depends on loop-extruding cohesin. (A) Experimental setup used in panels B-C and S1D-E (up), and corresponding Western blot showing Scc2-and Wpl1-depletion (down). (B) Smc6-FLAG enrichment in the centromeric region of chromosome 10, 420 – 450 kb from left telomere, in wild-type, Scc2-or Wpl1-depleted G2/M cells, as determined by ChIP-seq. The Y-axis shows fold enrichment of ChIP / input in linear scale, the X-axis shows chromosomal positions. Blue horizontal bars in the uppermost genomic region panel denote open reading frames (ORFs), CEN core centromere, ARS replication origins, and TER replication termination sites. (C) Smc6-FLAG enrichment at centromere 9 and in intergenic regions between indicated convergent gene pairs, and within the *BPH1* ORF, a Smc5/6-“non-binding site”, as determined by ChIP-qPCR analysis of wild-type, Scc2-or Wpl1-depleted G2/M cells. *N=3,* n.s.*: p>0.05*, *_*_ : p≤0.05, _**_ : p≤0.01, _***_ : p≤0.001*. (D) Experimental setup used in panels E-G (up), and corresponding Western blot showing Scc2-and Wpl1-depletion (down). (E) Smc6-FLAG enrichment along the arm of chromosome 7, 100 – 200 kb from left telomere, in wild-type and *top2-4* mutated G2/M-arrested cells, with or without Scc2-or Wpl1-depletion. Annotations as in panel (B). (F) Averaged Smc6-FLAG enrichment at Smc5/6 binding sites based on the analysis in (E). (G) Smc6-FLAG enrichment in intergenic regions between indicated convergent gene pairs as determined by ChIP-qPCR analysis of G2/M-arrested wild-type and *top2-4* mutated cells, with or without Scc2-or Wpl1-depletion. *N=3,* n.s.*: p>0.05*, *_*_ : p≤0.05, _**_ : p≤0.01, _***_ : p≤0.001*.

Smc5/6 accumulates at cohesin sites along chromosomes arms in response to the increase of SCIs caused by DNA replication in the absence of Top2 function (Jeppsson *et al*., 2014; Kegel *et al*., 2011). To test if this association also is under the control of loop-extruding cohesin, Scc2 or Wpl1 were depleted in G2/M-arrested *top2-4* cells grown at non-permissive temperature during the preceding S-phase (Figures 1D and S1F). Like the association around centromeres in wild-type cells, Scc2 depletion significantly reduced Smc5/6 chromosomal association in *top2-4* cells (Figure 1E-G). Moreover, depletion of Wpl1 caused an accumulation of Smc5/6 to levels even higher than after Top2 inhibition alone (Figure 1E-G). This shows that Smc5/6 binding to entangled chromosomes also requires loop-extruding cohesin.

### Smc5/6 chromosomal association requires ongoing transcription

The association of Smc5/6 to intergenic regions in between convergently oriented genes suggests that the complex might be regulated by transcription. We therefore investigated Smc5/6 chromosomal binding pattern after 3-, 15-, and 30-minutes treatment with the transcriptional inhibitor thiolutin (Figure 2A). Control experiments show that this causes RNA polymerase 2 (RNA pol II) to dissociate from chromosomes (Figures 2B and S2A). Thiolutin treatment caused a rapid and significant reduction of Smc6 around centromeres in in G2/M-arrested wild-type cells (Figures 2C, D, and S3A-B). Transcription inhibition also quickly and efficiently removed Smc5/6 from cohesin sites at centromeric regions and along chromosome arms in *top2-4* cells (Figures 2D-G, and S3A, C-D). Moreover, even if initially low, Smc5/6 association in between convergently oriented genes along chromosome arms in wild-type cells was also rapidly reduced by thiolutin (Figures 2G, S3D-E). In contrast, cohesin’s chromosomal association was only marginally affected by transcription inhibition in G2/M-arrested wild-type or *top2-4* cells (Figure S2B and S3F-G), in line with previous findings (Garcia-Luis *et al*., 2019; Jeppsson *et al*., 2022). We also investigated Smc5/6 localization by ChIP-qPCR after inhibition of the temperature sensitive RNA pol II allele *rpb1-1*, which confirmed that Smc5/6 dissociates from chromosomes following transcription inhibition in wild-type and *top2-4* cells (Figure S3H). Together, the results indicate that Smc5/6’s chromosomal association requires ongoing transcription of convergently oriented genes, in addition to cohesin loop extrusion. Given that the transcribing RNA polymerases overwind the DNA ahead of them, the mechanism that recruits Smc5/6 to intergenic regions in between convergently oriented genes could involve positively supercoiled DNA. This said, the transcription-dependent chromosomal positioning could also reflect that RNA pol II creates a boundary for loop-extruding cohesin (Banigan *et al*., 2023; Jeppsson *et al*., 2022), which in turn regulates Smc5/6 binding.

**Figure 2.**
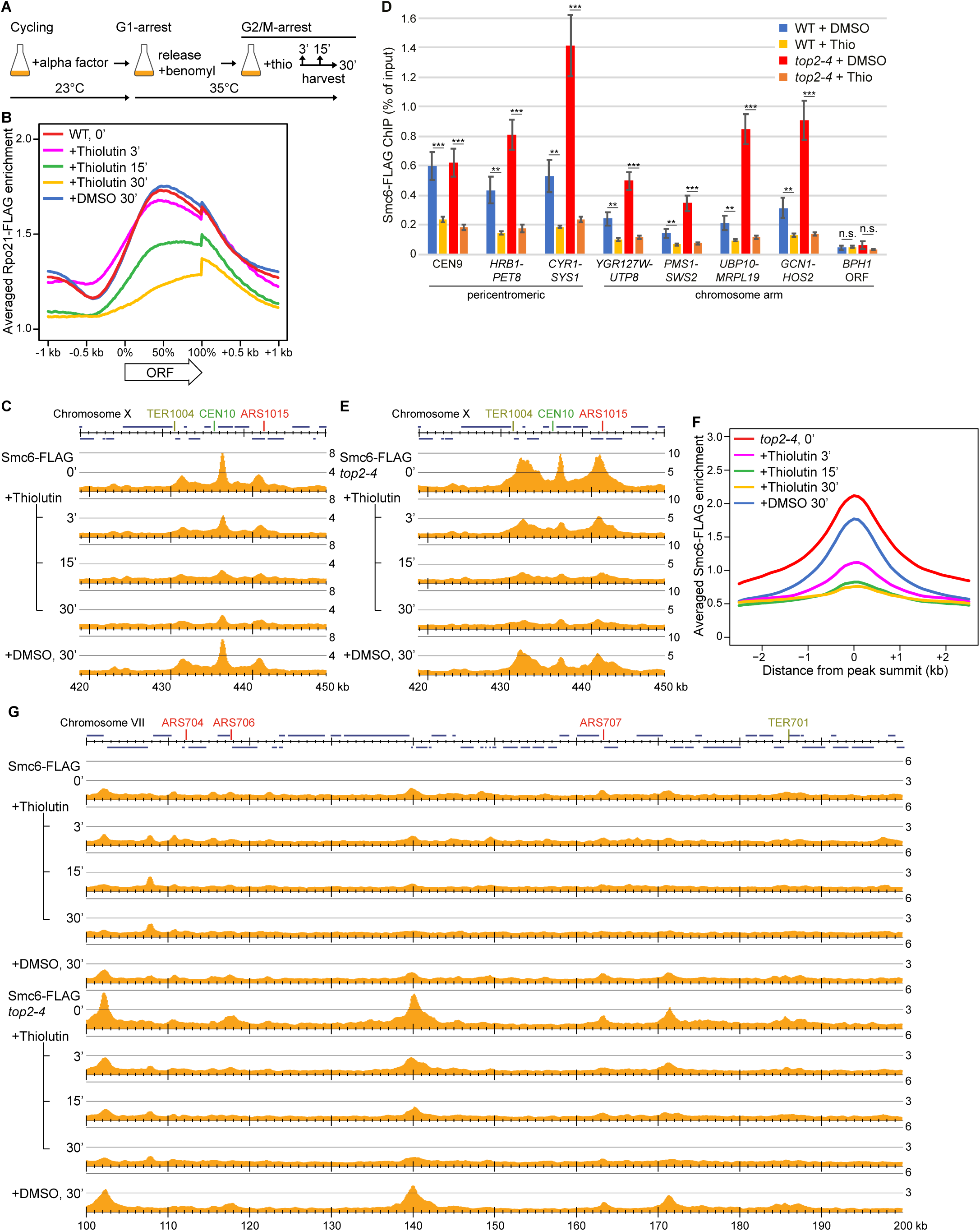
Smc5/6 chromosomal association requires ongoing transcription. (A) Experimental setup used for experiments displayed in panels B-G and Figures S2 and S3A-G. (B) Averaged Rpo21-FLAG enrichment within, and 1 kb up-and downstream ORFs, with or without preceding thiolutin-treatment during indicated periods. (C) Smc6-FLAG enrichment in the centromeric region of chromosome 10, 420 – 450 kb from left telomere, in wild-type cells, with or without indicated periods of thiolutin-treatment. Annotations as in Figure 1B. (D) Smc6-FLAG enrichment at centromere 9 and in intergenic regions between indicated convergent gene pairs, and within the *BPH1* ORF, a Smc5/6-“non-binding site”, as determined by ChIP-qPCR analysis of wild-type and *top2-4* cells, with or without preceding 30 minutes thiolutin-treatment. *N=3,* n.s.*: p>0.05*, *_*_ : p≤0.05, _**_ : p≤0.01, _***_ : p≤0.001.* (E) Smc6-FLAG enrichment as in (C), but in *top2-4* cells. (F) Averaged Smc6-FLAG enrichment at Smc5/6 binding sites based on the analysis in (G). (G) Smc6-FLAG enrichment along the arm of chromosome 7, 100 – 200 kb from left telomere, in wild-type (up) and *top2-4* mutated (down) G2/M-arrested cells, with or without preceding thiolutin-treatment during indicated periods. Annotations as in Figure 1B.

### Positive supercoiling recruits Smc5/6 to chromosomes

To conclusively assess if DNA supercoiling directly controls Smc5/6 chromosomal positioning independently of cohesin, we replaced the promoters of the *MCR1-DBR1* convergent gene pair with the strong constitutive promoters P_ADH1_ and P_GPD_, respectively (Figure 3A, S4A). This increased the transcription rates of *MCR1* ≈ 22 fold and *DBR1* ≈ 63 fold (Figure 3B), which will trigger an accumulation of positive supercoils between the genes (Garcia-Rubio and Aguilera, 2012). Aware of the role of Smc5/6 during S-phase, we ensured that the promoter replacement did not interfere with cell cycle progression or resistance to replication stress (Figure S4B-C). ChIP experiments showed that without promoter replacement some cohesin, but little, if any, Smc5/6 can be detected in between *MCR1* and *DBR1* (Figures 3C-D, S4A and S5A). In sharp contrast, high levels of convergent transcription caused a site-specific recruitment of Smc5/6 to the intergenic region, while cohesin enrichment was reduced (Figures 3C-D, and S5A-B). Moreover, the induced Smc5/6 association between the highly expressed convergent gene pair remained unchanged when cohesin function was inhibited using the temperature sensitive *scc1-73* allele (Figure 3E). Together, this indicates that Smc5/6 accumulation at this highly positively supercoiled chromosomal region occurs independently of cohesin. To test this further, we replaced the P_ADH1_ promoter with P_Gal1-10_ to enable inducible strong expression of *MCR1*, and thereby also cell cycle-specific induction of high convergent transcription (Figure 3F). Control experiments confirmed strong induction of *MCR1* upon galactose addition to the cell culture media (Figure S5C). Thereafter, cells were arrested in G1 or G2/M and convergent transcription was induced for 1h, after which Smc5/6 localization was investigated by ChIP-qPCR. This revealed that inducible convergent transcription also triggers Smc5/6 recruitment between the gene pair, not only in G2/M but also in a G1-arrest, when cells contain very low levels of cohesin, and Smc5/6 is normally not detected between convergent gene pairs (Figures 3G, S5D-F). The association of Smc5/6 to an un-replicated chromosome in G1-arrested cells confirms that Smc5/6 recruitment caused by strong convergent transcription can occur independently of cohesin. Importantly, it also shows that Smc5/6 can bind chromosomes independently of any other G2/M-specific factor, such as the presence of SCIs.

**Figure 3.**
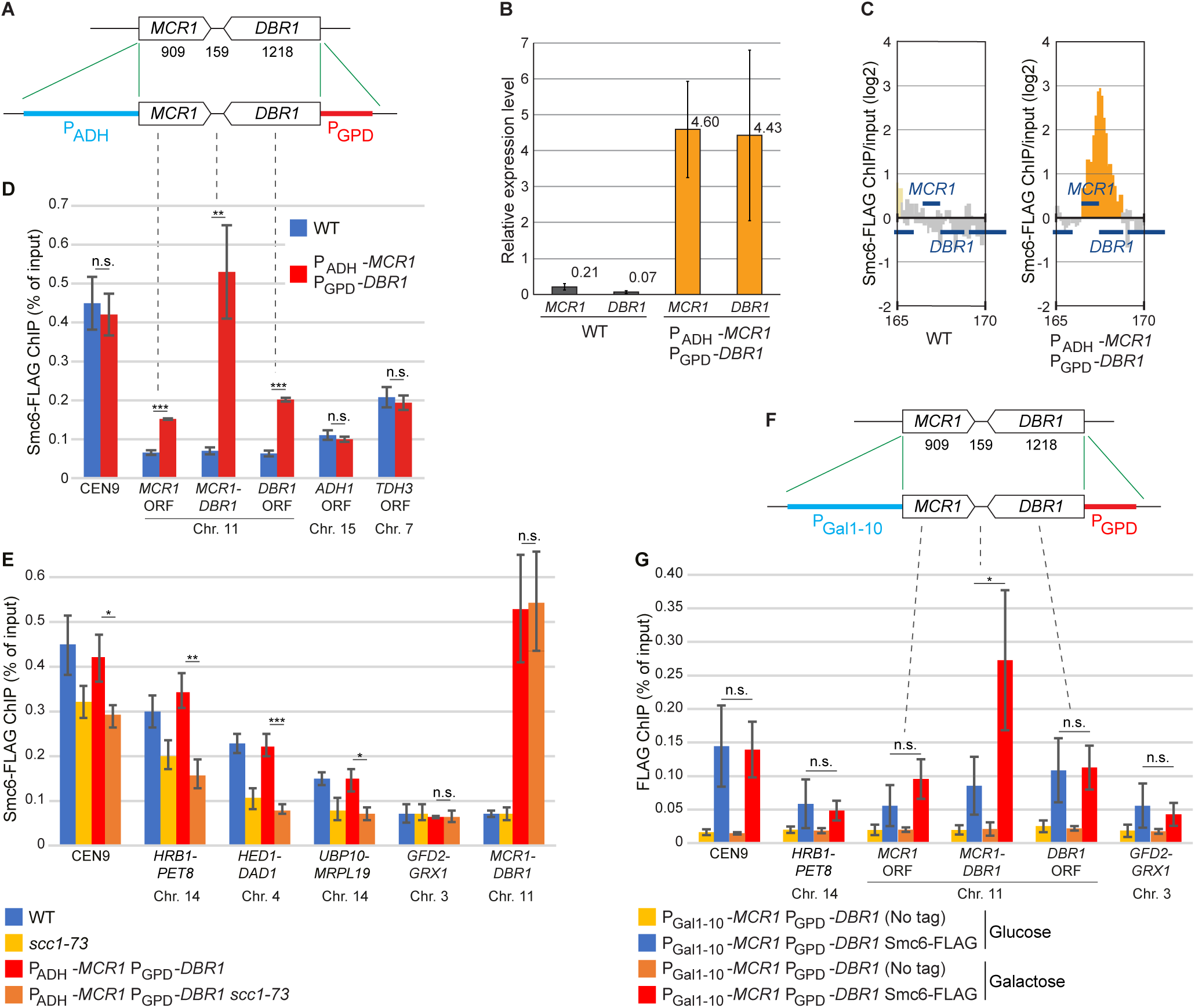
Positive supercoiling induced by site-specific high convergent transcription recruits Smc5/6 to chromosomes. (A) Genomic modifications used to create site-specific high convergent transcription. The size of *MCR1* and *DBR1* ORFs, and the intervening intergenic region, are indicated in number of base pairs. (B) Relative expression level of *MCR1* and *DBR1* as compared to *ACT1*, before and after promoter replacement, determined by RT-qPCR. *N=3,* error bars indicate standard deviations. (C) Smc6-FLAG enrichment between *MCR1* and *DBR1* before and after promoter replacement, determined by ChIP-on-chip. Blue vertical lines denote ORFs, the Y-axis shows fold enrichment of ChIP / input in log_2_ scale and the X-axis chromosomal positions on chromosome 11. (D) Smc6-FLAG enrichment at centromere 9, in the intergenic regions between the *MCR1-DBR1* convergent gene pair, and within *MCR1*, *DBR1*, *ADH1*, *TDH3* ORFs as indicated. The data was obtained by ChIP-qPCR analysis of cells with or without *MCR1* and *DBR1* promoter replacement. *N=3,* n.s.*: p>0.05*, *_**_ : p≤0.01, _***_ : p≤0.001.* (E) As in (D) but in wild-type or *scc1-73* temperature-sensitive mutant background cells, at centromere 9 and in between indicated convergent gene pairs, including the modified *MCR1*-*DBR1* site. (F) Genomic modifications used to create site-specific, galactose-inducible high convergent transcription. (G) Smc6-FLAG enrichment at centromere 9, at intergenic regions between indicated convergent gene pairs, and within *MCR1* and *DBR1* ORFs as indicated. The data was obtained by ChIP-qPCR analysis of G1-arrested cells with or without *MCR1* and *DBR1* promoter replacement, after 1 h in the presence of glucose or galactose. *N=3,* n.s.*: p>0.05*, *_*_ : p≤0.05*.

To investigate the effect of positive DNA supercoiling on Smc5/6 association to unmodified genomic sites, Top1 and Top2, both of which are required to resolve (positive) supercoils, were inactivated in G2/M-arrested cells, and Smc5/6 chromosomal association was investigated (Figure 4A and S6A). Inactivating Top2 after DNA replication alone does not increase the level of SCIs or lead to Smc5/6 accumulation along chromosome arms (Figure 4B-D) (Jeppsson *et al*., 2014). Smc5/6 binding was also unaltered after Top1 depletion alone. However, Smc5/6 enrichment between convergently oriented genes was significantly increased after inactivation of both topoisomerases, (Figures 4B-D, and S6A). In addition, the complex accumulated in broad peaks that spread into the flanking open reading frames (Figure 4B-C). This, together with the results obtained from the artificially highly expressed convergent gene pair, show that positive DNA supercoils determine Smc5/6 chromosomal positioning.

**Figure 4.**
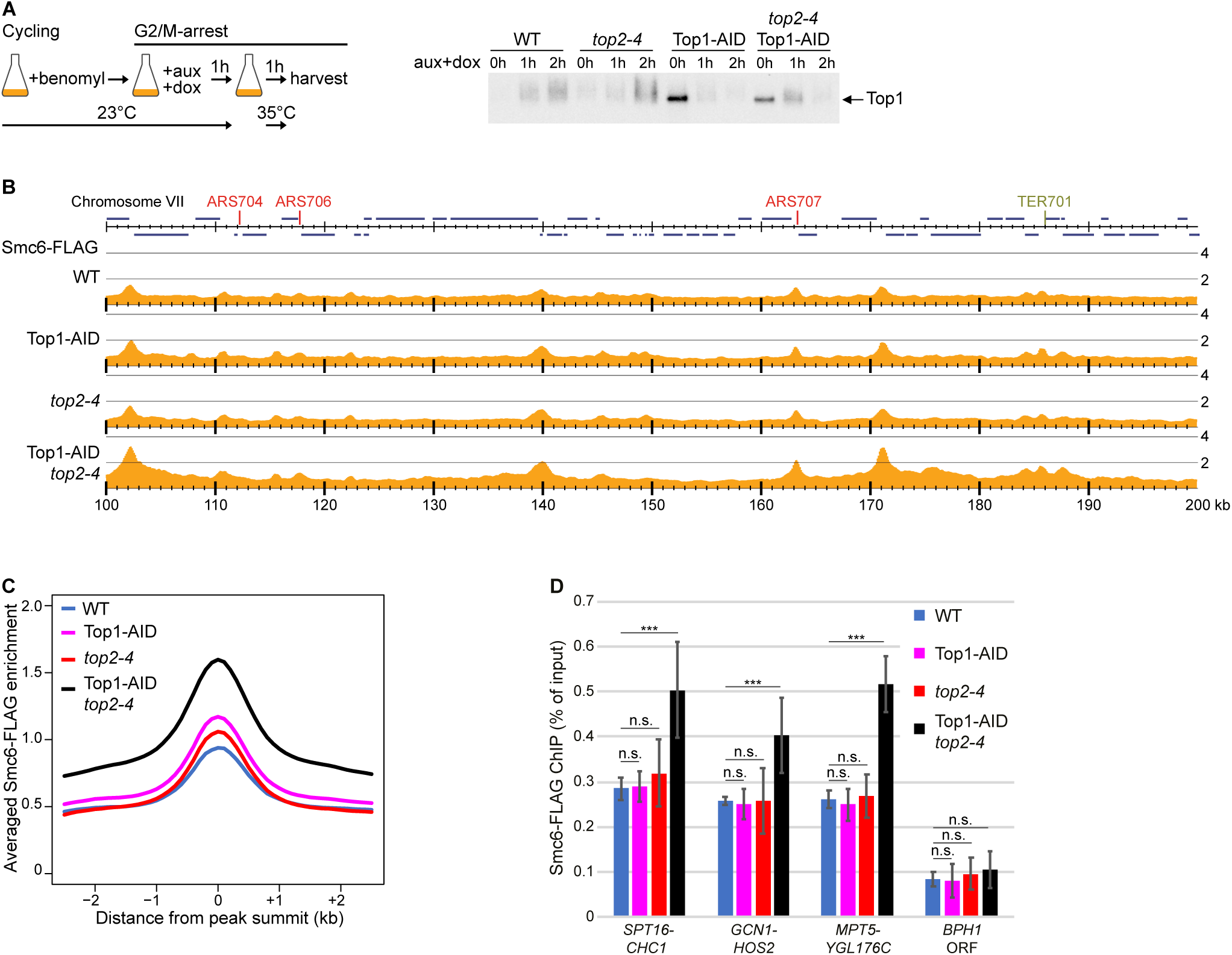
Positive supercoiling recruits Smc5/6 to endogenous chromosomal sites. (A) Experimental setup used in panels B-D and S6A (left), and corresponding Western blot showing Top1-depletion (right). (B) Smc6-FLAG enrichment along the arm of chromosome 7, 100 – 200 kb from left telomere, in G2/M-arrested cells, with or without preceding Top1-depletion, *top2-4* inhibition, or both. Annotations as in Figure 1B. (C) Averaged Smc6-FLAG enrichment at Smc5/6 binding sites based on analysis in (B). (D) Smc6-FLAG enrichment in intergenic regions between indicated convergent gene pairs, and within the *BPH1* ORF, a Smc5/6-“non-binding site”, as determined by ChIP-qPCR analysis of the same cell types as in B-C. *N=3,* n.s.*: p>0.05*, *_*_ : p≤0.05, _**_ : p≤0.01, _***_ : p≤0.001*.

#### Transcription levels and gene length of surrounding genes govern Smc5/6 enrichment

To gain further insight into the regulation of Smc5/6 chromosomal distribution and enrichment level, we also performed multivariable analysis by computational machine-learning using 47 defined chromosomal features (Table S4), which were organized into 8 groups according to their properties (Figure 5A). The analysis focused on all intergenic regions (IGRs) in the *S. cerevisiae* genome and was based on ChIP-seq of Smc5/6 and cohesin. When all combined, the defined features were sufficient to accurately predict the enrichment level of both Smc5/6 and cohesin at each IGR (Figure 5A). For cohesin, modeling solely based on group D features resulted in a good prediction (Figure 5A), which nicely reflects previous reports that cohesin enrichment mainly depends on the convergent orientation of surrounding genes (Lengronne *et al*., 2004). Addition of group H features, which describe chromosome length, and distance to centromeres and telomeres, had a significant positive impact on the predictive power of the analysis, likely reflecting the enrichment of cohesin in the centromeric area (Figure S7A-B, and Table S4) (Lengronne *et al*., 2004). In line with that Smc5/6 and cohesin show a high degree of colocalization on entangled chromosomes, the features describing convergent orientation of genes (feature group D), and positioning along chromosomes (feature group H) were also important to predict Smc5/6 chromosomal distribution in *top2-4* cells (Figure 5B-C). However, to accurately predict Smc5/6 enrichment in wild-type cells, the features describing transcription rates and length of the most proximal, adjoining convergent gene pair (feature group B) had to be included (Figures 5B-C). Investigation of each feature revealed that Smc5/6 enrichment was associated with higher expression and longer length of the adjacent genes (Table S4). Knowing that high transcription and longer genes induce more positive supercoiling (Guo *et al*., 2021; Joshi *et al*., 2012), this strengthens the notion that Smc5/6 is controlled directly by positive DNA supercoiling.

**Figure 5.**
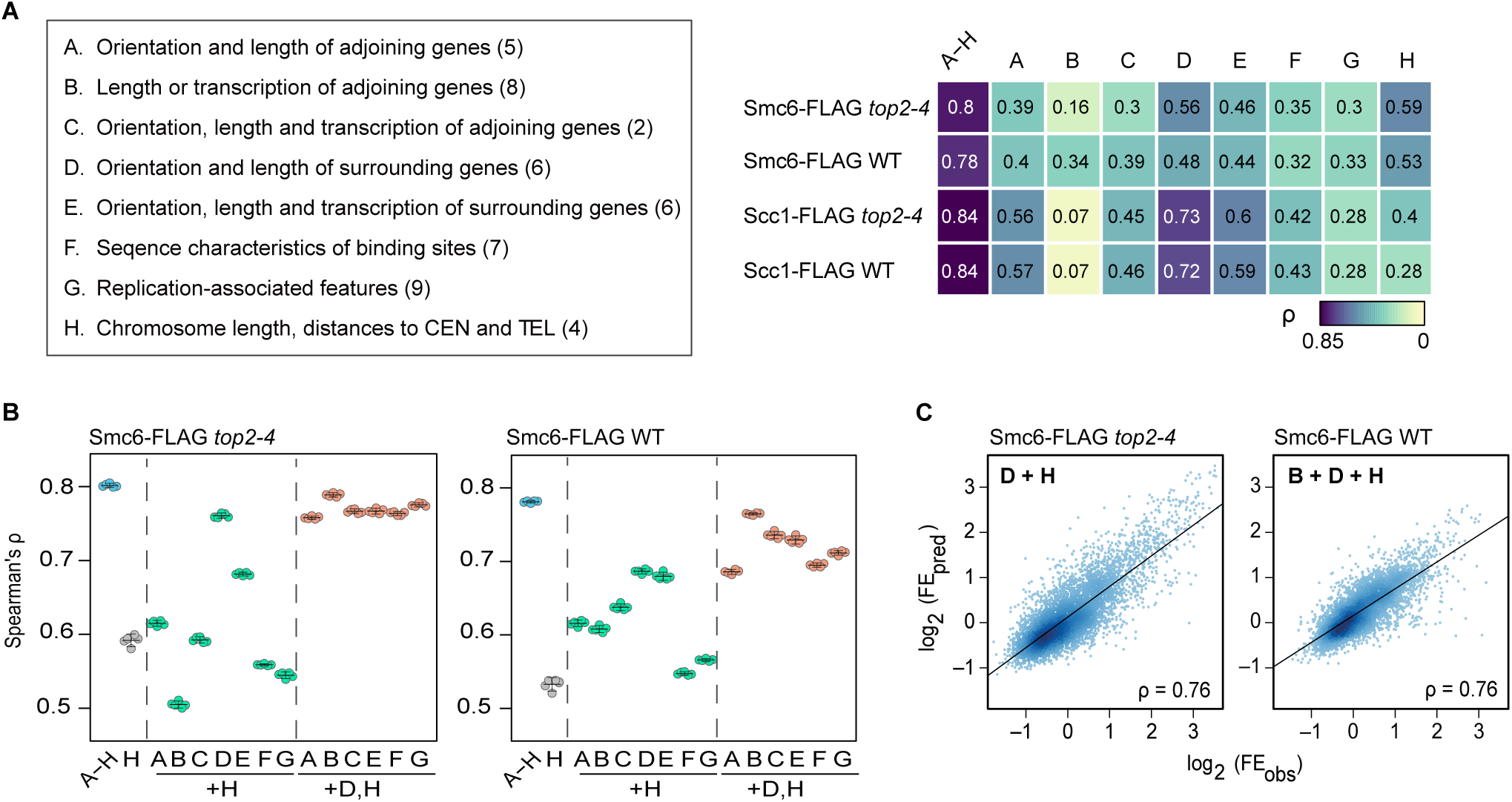
Transcription levels and gene length of surrounding genes govern Smc5/6 enrichment. (A) Left: Groups of features used for modeling (see Table S4 for details). The number of features per group are indicated in parentheses. Right: The fold-enrichment (FE) of Smc6 and Scc1 at each intergenic region in wild-type and *top2-4* mutated cells was modeled by machine learning. Groups of features used for model building are indicated at the top. Accuracy of the prediction was evaluated by Spearman’s rank correlation coefficient (ρ) between predicted and observed FE values. (B) Accuracy of prediction models based on indicated groups of features. Smc6-FLAG enrichment in *top2-4* and wild-type cells was predicted, and the prediction accuracy was evaluated as in (A). Four-fold cross-validation was repeated five times with differently partitioned datasets, and the resulting correlation coefficients are plotted. The horizontal line indicates the mean, and whiskers show the 95% confidence interval. (C) Representative prediction results of analysis in (B). Observed and predicted FE values, FE_obs_ and FE_pred_, respectively, were plotted in log_2_ scale. Feature groups D and H were used to model Smc6-FLAG enrichment in *top2-4* cells, and groups B, D and H for the modeling of Smc6-FLAG enrichment in wild-type cells.

### Smc5/6 initiates loop extrusion at the tip of positively supercoiled DNA plectonemes

To obtain mechanistic understanding of Smc5/6 function and association to positively supercoiled DNA, we performed single molecule analysis that allows direct visualization of Smc5/6 loop extrusion on supercoiled DNA attached to a passivated glass surface. Previously, we used this system to show that purified Smc5/6 can extrude DNA loops on nicked DNA as dimers of hexameric complexes (Smc5, Smc6 and Nse1-Nse4), while monomeric complexes translocate along DNA (Pradhan *et al*., 2023). The Nse5 and Nse6 subcomplex (Nse5/6) promotes translocation (and reduces looping) of Smc5/6 by inhibiting dimerization of Smc5/6 hexamers. We therefore purified hexameric and octameric Smc5/6 complexes, *i.e.*, with and without Nse5/6, and determined their association and loop extrusion activity on positively and negatively supercoiled 42 kb DNA substrates, stained by the intercalating SYTOX Orange (SxO) dye. Positive supercoiling was induced by adding SxO after restraining torsion release by tethering the DNA ends to the surface, and negative supercoils were created by reducing dye concentration after attaching pre-stained DNA, as previously reported (Kim *et al*., 2018). By using half of the SxO concentration that allowed maximum intercalation (C_1/2_ = 300 nM) for generation and visualization of positive and negative supercoils, similar levels of dynamic plectonemes, observed as transient local maxima in DNA kymographs, were obtained (Figures S8A-C).

Upon addition of hexameric Smc5/6 to positively supercoiled DNA, the dynamic plectonemes were rapidly gathered into a single stable loop by extrusion (Figures 6A-C, S8D and Movie 1). Strikingly, loop extrusion was 6.7±2 times more likely to occur on positively supercoiled as compared to nicked (Figure 6D). In contrast, loop extrusion initiation on negatively supercoiled DNA was not enhanced as compared to that observed on nicked DNA (Figure 6E). The rate of loop extrusion on positively supercoiled DNA was modestly increased to 2.4±1.3 kbp per sec, as compared to 1.8±1.1 kbp per sec on nicked DNA (Figure S8E), although the difference was not statistically significant. This reveals that positive, but not negative, DNA supercoiling greatly enhances Smc5/6 loop extrusion initiation rate.

**Figure 6.**
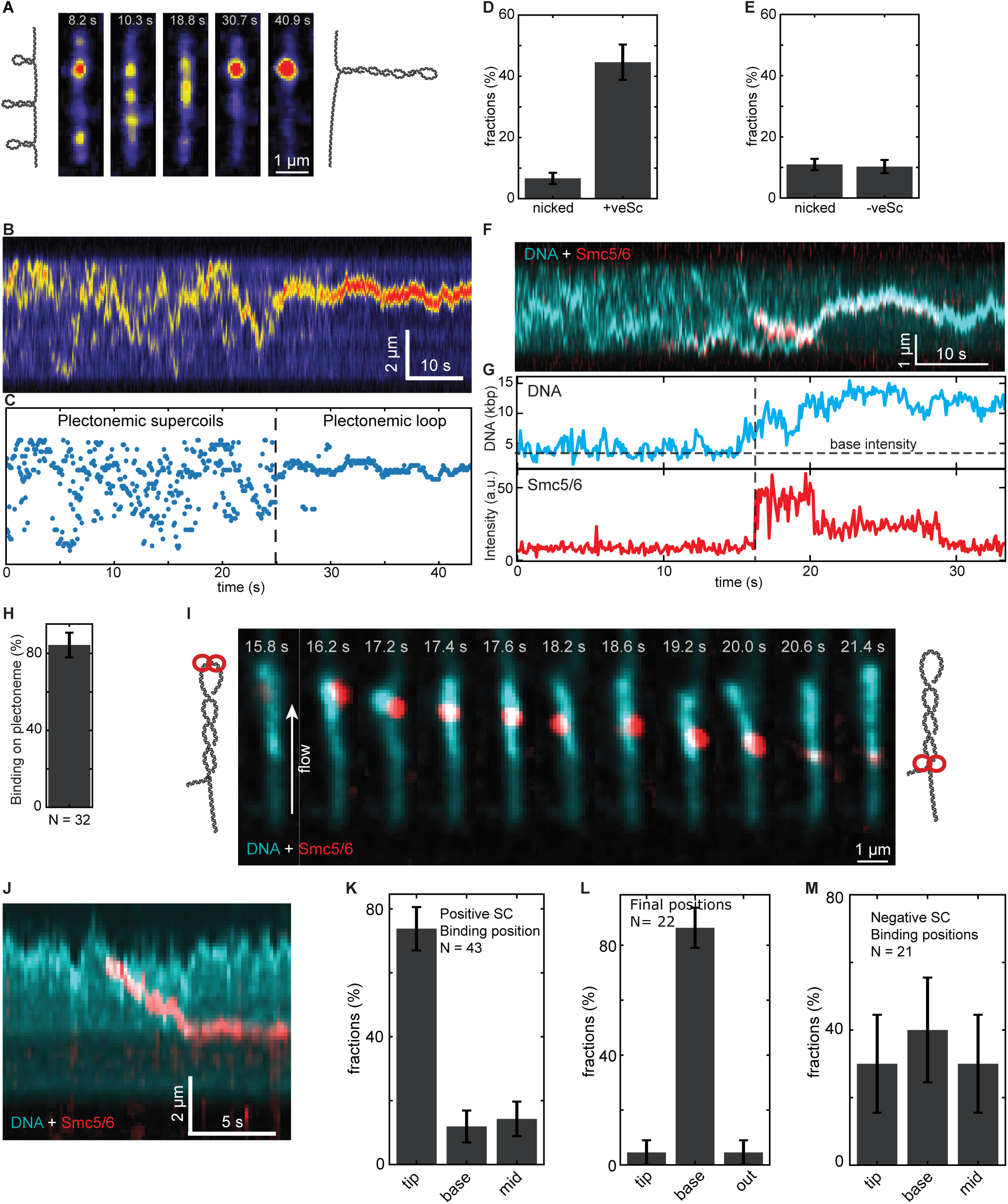
Smc5/6 preferentially binds and initiates DNA loop extrusion at the tip of positively supercoiled plectonemes. (A) Snapshots of positively supercoiled DNA stained with SYTOX Orange after the addition of 1 nM hexameric Smc5/6 in the presence of ATP. (B) A kymograph of the DNA, and (C) puncta positions, corresponding to panel (A). (D-E) The fraction of extruded loops by Smc5/6 per DNA molecule on positively (D) and negatively (E) supercoiled DNA, compared to nicked DNA. (F) A kymograph of DNA (cyan) loop extrusion by labeled hexameric Smc5/6 (red) on positively supercoiled DNA, initially binding at a plectoneme and resulting in a single large plectonemic loop. (G) Time trace of the DNA loop size and intensity of labeled Smc5/6 in (F). (H) The fraction of labeled hexameric Smc5/6 binding to plectonemes, with the remaining fraction binding outside the plectonemes. (I-J) Binding of labeled hexameric Smc5/6 (red) to a positively supercoiled plectoneme and following DNA (cyan) loop extrusion under buffer side-flow (I), and the corresponding kymograph (J). (K-L) Initial (K) and final (L) binding positions of labeled Smc5/6 on positively supercoiled DNA plectonemes under side-flow. M) Binding positions of labeled hexameric Smc5/6 on negatively supercoiled DNA.

Interestingly, analysis of fluorescently labeled hexameric Smc5/6 revealed that Smc5/6 initially binds to the dynamic positively supercoiled plectonemes, and following the swift depletion of the smaller dynamic plectonemes, Smc5/6 overlaps with a single intensifying spot (Figure 6F-H, Movie 2). Bleaching dynamics of the labeled Smc5/6 after initiation of loop extrusion revealed a larger fraction (58±8%) of two-step bleaching events as compared to single-step events (34±7%), which taken together with the labelling efficiency of 72%, confirms that Smc5/6 performs loop extrusion on positive supercoiled DNA as dimers of hexameric complexes, similarly to the extrusion detected on nicked DNA (Figures 6F-G, S8F-H and (Pradhan *et al*., 2023)).

Analysis using fluorescently labelled octameric Smc5/6, which primarily exhibits translocation in monomeric form, and very low levels of DNA loop extrusion (Pradhan *et al*., 2023), revealed that the binding of this version of Smc5/6 on supercoiled DNA was approximately 4 times higher as compared to nicked DNA (Figure S8I-J). Hence, the results show that preferential binding on positively supercoiled DNA is shared between the monomeric and dimeric forms of Smc5/6, and indicates that the enhanced looping probability on positively supercoiled DNA is due to increased binding of Smc5/6 dimers to positively supercoiled plectonemes.

Applying side-way buffer flow to the chamber pins the dynamic positive plectonemes into one single plectoneme (Kim *et al*., 2022), thus allowing observation of binding and the subsequent loop extrusion dynamics of the labeled Smc5/6 dimers along plectonemes. This revealed that the Smc5/6 dimer initially associates to the tip of the positively supercoiled plectoneme, and subsequently moves downwards until it is found at the base of the loop (Figure 6I-L and Movie 3). The stalling at the base of the loop indicates that the downward movement reflects loop extrusion, and accordingly, using the octameric complex which primarily exhibit translocation along DNA (Pradhan *et al*., 2023), stalling was not detected but instead Smc5/6 continued to the end of the tethered DNA molecule (Fig S7I and K). Moreover, in contrast to the preferred association to the tip of positively supercoiled plectonemes, Smc5/6 binding on flow-stretched negatively supercoiled plectonemes occurred at random positions (Figure 6M).

Altogether, the results remarkably show that dimers of Smc5/6 specifically recognize the tip of positively supercoiled DNA plectonemes, and efficiently initiates loop extrusion to gather the surrounding supercoiled DNA into one large plectonemic loop.

### Smc5/6 links positively supercoiled chromosomal loci

The fact that Smc5/6 preferentially initiates loop extrusion on the tip of positively supercoiled DNA *in vitro* (Figure 6), opens for that Smc5/6’s association to transcription-induced positive supercoiled chromosomal loci (Figures 1-4) reflects that Smc5/6 performs loop extrusion at these sites *in vivo*. To explore this hypothesis, Hi-C analysis was performed in G2/M-arrested cells that were depleted for Smc5 and Smc6 from the preceding G1-phase (Figures 7A and S9A-B). Supporting the idea that Smc5/6 performs DNA loop extrusion *in vivo*, the Hi-C analysis revealed a small, but significant and reproducible reduction of short-range (around 10-80 kb) chromosomal *cis* interactions in the Smc5/6-depleted cells (Figure 7B-E). In contrast, Smc5/6 does not contribute to cohesin-mediated loops since they remained intact after the depletion (Figure 7F).

**Figure 7.**
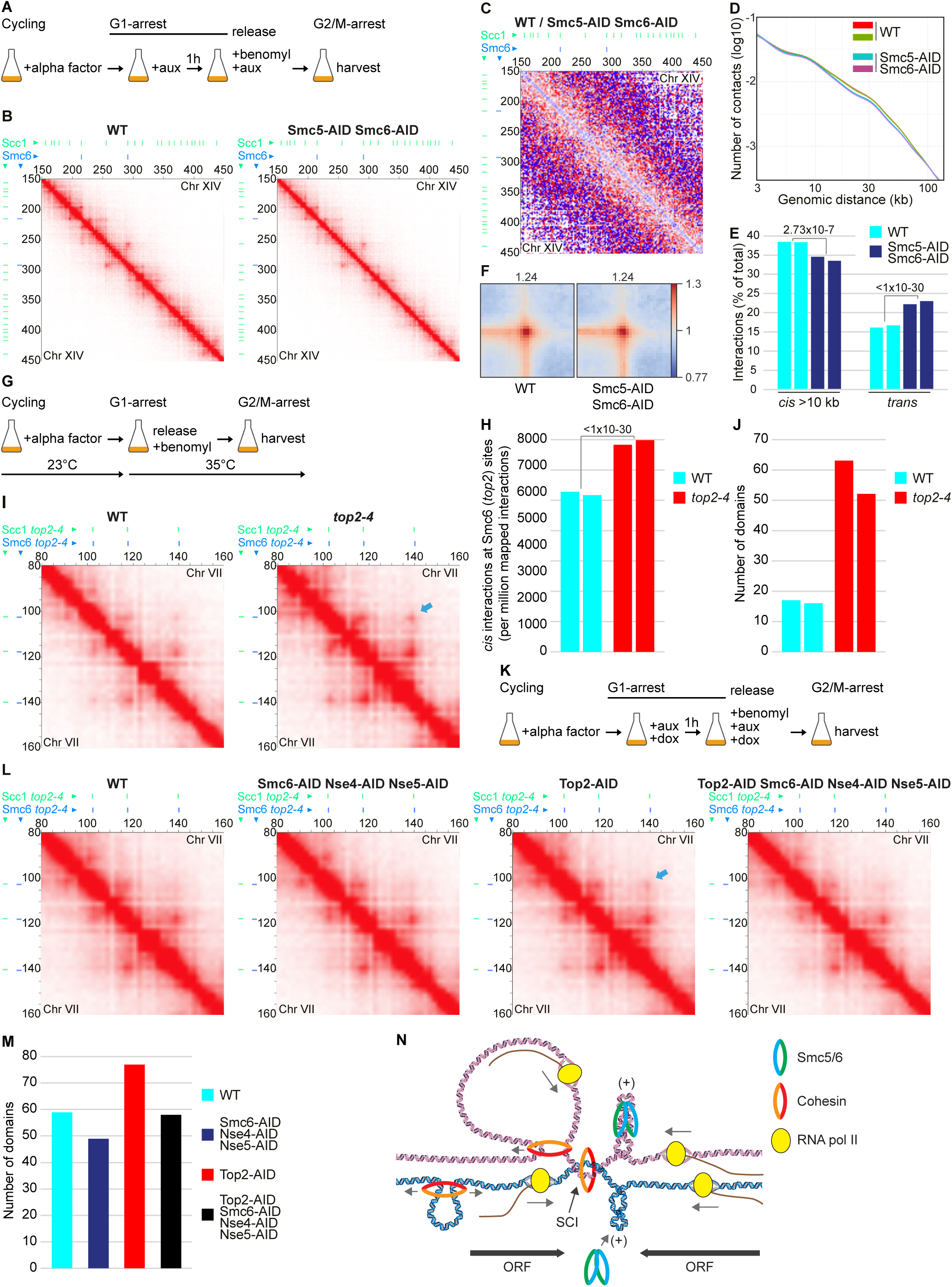
Smc5/6 links positively supercoiled chromosomal loci. (A) Experimental setup used in panels B-F. (B) Normalized Hi-C contact maps (2 kb binning) showing *cis* interactions along the arm of chromosome 14, 150 - 450 kb from left telomere, in G2/M-arrested wild-type and Smc5/6-depleted (Smc5-AID Smc6-AID) cells. Lines on top and to the left of the panels: green = cohesin binding sites, blue = Smc5/6 binding sites. (C) Normalized Hi-C ratio maps (2 kb binning) comparing chromosome *cis* interactions in G2/M-arrested wild-type and Smc5/6-depleted cells along the same chromosomal regions as depicted in B). (D) Contact probability plots as function of genomic distance comparing interactions in G2/M-arrested wild-type and Smc5/6-depleted cells. (E) Quantification of *cis* (>10 kb) and *trans* interactions in G2/M-arrested wild-type and Smc5/6-depleted cells. (F) Pile-up plots of averaged *cis* interactions in wild-type and Smc5/6-depleted cells between all pairs of cohesin sites situated 20 - 40 kb apart on chromosome arms. (G) Experimental setup used in panels H-J, S9D and S10A. (H) Quantification of *cis* interactions (>10 kb) in wild-type and *top2-4* cells, anchored at Smc5/6 chromosome arm binding sites in *top2-4* cells. (I) Normalized Hi-C contact maps (2 kb binning) showing *cis* interactions along the arm of chromosome 7, 80 - 160 kb from left telomere, in G2/M-arrested wild-type and *top2-4* cells. Lines on top and to the left of the panels: green = cohesin binding sites in *top2-4* mutant, blue = Smc5/6 binding sites in *top2-4* mutant, light blue arrow: example of increased *cis* interactions between Smc5/6 binding sites in *top2-4* cells. (J) Number of domains in G2/M-arrested wild-type and *top2-4* cells. (K) Schematic description of the experimental setup used in panels L-M and S10D-E. (L) Normalized Hi-C contact maps (2 kb binning) showing *cis* interactions along the arm of chromosome 7, 80 - 160 kb from left telomere, in wild-type, Smc6-AID Nse4-AID Nse5-AID triple depletion, Top2-AID, and Top2-AID Smc6-AID Nse4-AID Nse5-AID quadruple depletion cells. Lines on top and to the left of the panels: green = cohesin binding sites in *top2-4* mutant, blue = Smc5/6 binding sites in *top2-4* mutant, light blue arrow: example of increased Smc5/6-dependent *cis* interactions between Smc5/6 binding sites in Top2-AID cells. (M) Number of domains in G2/M-arrested wild-type, Smc6-AID Nse4-AID Nse5-AID triple depletion, Top2-AID, and Top2-AID Smc6-AID Nse4-AID Nse5-AID quadruple depletion cells. (N) A model summarizing that cohesin loop extrusion, transcription, gene orientation and chromosome entanglements (SCIs) contribute to the formation of positively supercoiled DNA plectonemes, which are recognized by dimers of hexameric Smc5/6 that initiates loop extrusion.

Our results suggest that the accumulation of Smc5/6 on transcription-induced positive supercoiled DNA can be further strengthened by the presence of SCIs (Figure 2). If Smc5/6 performs loop extrusion at these sites, SCIs are expected to increase the level of chromosome *cis* interactions. To test this idea, Hi-C analysis was performed on G2/M-arrested cells which had replicated in the absence of functional Top2 during the preceding S-phase (Figures 7G and S9C). The Hi-C results revealed a significant increase of *cis* interactions at Smc5/6 binding sites that appear along chromosome arms after Top2 inhibition (Figure 7H). The increased *cis* interactions in *top2-4* occasionally linked neighboring Smc5/6 binding sites, indicative of a loop forming between them (Figures 7I and S9D-F). However, overall, the averaged strength of loops was not increased in *top2-4* cells (Figure S10A). Instead, the number of domains was strongly increased following Top2 inactivation (Figure 7J). These findings are in line with the idea that Smc5/6 performs loop extrusion after recruitment to positive supercoils on the entangled chromosomes, although rarely forming stable loops between neighboring detected binding sites. In further support of this, simultaneous depletion of Smc5/6 (Smc6-AID, Nse4-AID Nse5-AID) and Top2 (Figure 7K and S10B-C), abolished the increase of *cis* interactions at Smc5/6 binding sites (Figures 7L and S10D-E), and reduced the number of domains back to wild-type levels (Figure 7M). Together this supports a model where Smc5/6 recognizes the tip of transcription-induced positive supercoiled DNA plectonemes that arise at cohesin loop boundaries, and initiates DNA loop extrusion (7N).

## DISCUSSION

Here, we show that Smc5/6 is recruited to transcription-induced positive DNA supercoils that appear at the base of cohesin-dependent chromosomes loops in the genome, and links positively supercoiled chromosomal loci in *cis*. Single-molecule imaging analysis provides detailed mechanistic insight into how Smc5/6 performs this function by revealing that dimers of Smc5/6 preferentially binds and initiates DNA loop extrusion at the tip of positively supercoiled plectonemes. This discovery of a chromosome organization function for Smc5/6 unites this so far enigmatic complex with cohesin and condensin, which also control the spatial organization of chromosomes using DNA loop extrusion.

The starting point of our investigation was the earlier observations that Smc5/6 colocalizes with cohesin between convergently oriented genes and is connected to DNA supercoiling (Gutierrez-Escribano *et al*., 2020; Jeppsson *et al*., 2014; Kegel *et al*., 2011; Serrano *et al*., 2020). This suggested that the complex could be jointly controlled by cohesin and transcription-induced supercoiling, and accordingly, we found that Smc5/6 association is controlled by Scc2, Wpl1 and ongoing transcription (Figures 1 and 2). To tease out the role of transcription-induced DNA supercoiling in Smc5/6 chromosomal association, we created the genetically engineered convergent gene pair, in which both genes are transcribed at exceptionally high levels, thereby creating strong positive supercoiling in the intergenic region (Figure 3). The finding that Smc5/6 is site-specifically recruited in between the two highly expressed convergently oriented genes independently of cohesin (Figure 3), and the increase of Smc5/6 at endogenous binding sites after concomitant depletion of Top2 and Top1(Figure 4) pinpoint positive supercoiling as the main underlying factor controlling the chromosomal binding pattern of the complex. Furthermore, single molecule analysis establishes that Smc5/6 recognizes and preferentially binds the tip of positively supercoiled plectonemes to efficiently initiate loop extrusion, finally confirming positive supercoiled DNA plectonemes as preferred substrates for Smc5/6 (Figure 6).

Our observations that supercoils accumulate at the base on cohesin loops in the yeast genome gain support from investigations in mammalian cells showing that Top2 is active at TAD boundaries, and that this activity is controlled by cohesin, the TAD boundary protein CTCF, and transcription (Canela *et al*., 2017; Gittens *et al*., 2019; Uuskula-Reimand *et al*., 2016). Human Smc5/6 chromosomal association also follows that of cohesin, appearing on chromosomes before replication and remaining until mitosis when both complexes concentrate in the centromeric area (Venegas *et al*., 2020; Wendt *et al*., 2008). Furthermore, Smc5/6 chromosomal enrichment was also recently shown to be reduced upon transcription inhibition (Aurélie *et al*., preprint on bioRxiv: https://doi.org/10.1101/2023.05.04.539344), well in line with a highly conserved regulation of the complex in human and yeast cells.

Comparison of Smc5/6 binding patterns with that of the bacterial protein GAPR, which binds positive supercoiling in the form of overtwisted DNA (Guo *et al*., 2018), also reveals interesting similarities. When expressed in yeast, GAPR is found at the 3’ end of most genes, is enriched in between convergently oriented genes, and positively correlates with transcription strength (Guo *et al*., 2021). Consequently, GAPR is also found at most cohesin binding sites, including those in the pericentromeric region. These observations support the notion that Smc5/6 is found in regions where chromosomal DNA is positively supercoiled. In addition, GAPR binding pattern indicates that positive supercoiling can be generated from upstream co-oriented genes, and similarly, Smc5/6 positioning and enrichment are not only influenced by the most proximal gene pair, but also by the orientation of genes up to 10 kb away from the binding sites (feature group D, Figure 5 and Table S4). This said, the preferential association of Smc5/6 to certain cohesin sites, in comparison to the more wide-spread distribution of GAPR, suggests that

Smc5/6 associates with a specific feature of positive supercoiling. Based on our observation that Smc5/6 preferentially recognizes the tip of positively supercoiled plectonemes (Figure 6), Smc5/6 potentially indicates the positions where these fundamental, but so far elusive, chromosomal structures are formed or stabilized.

Since plectoneme formation requires a certain threshold of overtwisting, Smc5/6 is expected to be enriched in regions of high levels of positive supercoiling. Accordingly, exceptionally strong convergent transcription triggers the association of Smc5/6, which also can occur in G1-arrested cells and in the absence of cohesin (Figure 3). In addition, Smc5/6 should be detected at sites where other features facilitate the transition of overtwisting into plectonemes. One such feature would be factors that confine the superhelical twist, *i.e.*, prevent it from spreading over larger chromosomal regions which reduces the local level of twist. Interestingly, analysis of the effects of supercoiling on transcription in budding yeast suggests that supercoils are more confined on longer chromosomes and far away from chromosome ends, which mimics Smc5/6 distribution in wild-type cells (Jeppsson *et al*., 2014; Joshi *et al*., 2010; Kegel *et al*., 2011). Similarly, Smc5/6 accumulation at the base on cohesin loops and along intertwined chromatids suggest that cohesin confines transcription-induced supercoiling at the base of the loops, and that this confinement is strengthened by SCIs.

Another question emanating from our investigation is how Smc5/6 recognizes the tip of positively supercoiled plectonemes specifically. This will of course demand detailed structural analysis to reveal, but it is interesting to notice that also condensin preferentially binds and loop extrudes positively supercoiled DNA (Kim *et al*., 2022). This said, while Smc5/6 performs two-sided loop extrusion in the form a dimer of complexes, condensin loop extrusion is one-sided and is executed by monomeric complexes. The rate of condensin loop extrusion was also lowered by supercoiling, which was not observed for Smc5/6. Thus, even if the initial recognition of the positively supercoiled DNA might be similar, the following reaction will differ. Our Hi-C analysis suggests that after loading, Smc5/6 uses loop extrusion to reel in DNA (Figure 7), which occasionally can bring two Smc5/6 binding sites together in the three-dimensional space. This said, the finding that chromosome intertwinings leads to an Smc5/6-dependent increase in domains, but does not affect loops, suggests that the extrusion process is most often disrupted before reaching all the way to the neighboring Smc5/6 binding sites. The fact that Smc5/6 promotes segregation of entangled chromosomes (Jeppsson *et al*., 2014), might indicate that this extrusion activity supports chromosome segregation and viability, an interesting focus for future studies.

Another interesting outcome of the initial Smc5/6 loading at the tip of positive plectonemes can be proposed based on a recent study suggesting that Smc5/6 relies on positive supercoiling when acting as a viral restriction factor (Aurélie *et al*., preprint on bioRxiv: https://doi.org/10.1101/2023.05.04.539344). This function leads to transcriptional silencing on small circular episomal DNA, which has been suggested to depend on topological entrapment of DNA within the Smc5/6 complex (Abdul *et al*., 2022; Kanno *et al*., 2015; Taschner *et al*., 2021). Based on this, our finding that Smc5/6 specifically recognizes the tip of positively supercoiled plectonemes (Figure 6) might disclose the mechanism by which the complex initiates topological entrapment and viral restriction. After the initial binding at positive supercoils, Smc5/6 could either entrap DNA immediately, or initiate loop extrusion or translocation, to later be converted into the topological entrapment binding-mode.

All taken together, this investigation uncovers Smc5/6’s function *in vivo*, and explains the role of the complex’s recently discovered loop extrusion activity. Crucially, the analysis also shows that transcription-induced positive supercoiling is not only a problem for topoisomerases to resolve, but also acts as a central regulator of three-dimensional chromosome organization via Smc5/6.

## ACKNOWLEDGEMENT

We thank C. Dekker and J. van der Torre for sharing the plasmids and the protocol for making 42kb coilable DNA construct. The study was funded by JSPS Postdoctoral Fellowship for Overseas Researchers, FY2016, (to KJ), Max Planck Society and European Research Council Starting Grant no. 101076914 (to EK), JST CREST Grant Number JPMJCR18S5 and JSPS KAKENHI Grant Numbers JP20H05686, JP20H05940 (to KS), JSPS KAKENHI Grant Number JP21K06012 (to TSu), and Swedish Cancer foundation, Swedish research council, and Centre for Innovative Medicine (CIMED) (to CB).

## AUTHOR CONTRIBUTION

KJ and CB conceived the initial ideas, and developed the study together with all co-authors. KJ performed all ChIP, RNA-seq and Hi-C experiments except for the ChIP-qPCR experiments presented in 4D and 3E, 3G, S5E that were executed by MUI and DGB, respectively. TK purified and labelled the hexameric Smc5/6 for single molecule analysis. BP performed the single molecule experiments and data analysis, TaSu executed the machine learning analysis, ToSa, RN, KJ and KS performed and developed the bioinformatic analysis of ChIP and Hi-C analyses. EK supervised the single molecule imaging part of the project. KJ and CB wrote the manuscript with input from all authors. KJ, KS, EK, and CB acquired funding.

## METHODS

### RESOURCE AVAILABILITY

#### Lead contact

Further information and requests for resources and reagents should be directed to and will be fulfilled by the lead contact, Kristian Jeppsson.

#### Materials availability

Yeast strains generated for this study are available upon request to the lead contact.

#### Data and code availability

ChIP-seq, RNA-seq and Hi-C data will by publicly available as of the date of publication. Any additional information required to reanalyze the data reported in this paper is available from the lead contact upon request.

## EXPERIMENTAL MODEL

The *in vivo* experiments were performed in budding yeast *S. cerevisiae* of W303 origin (*ade2-1 trp1-1 can1-100 leu2-3,112 his3-11,15 ura3-1 RAD5*) with the modifications listed in Table S1. Purified hexameric and octameric *S. cerevisiae* Smc5/6 complexes were used in the single molecule analysis.

## METHOD DETAILS

### Yeast growth conditions, protein degradation and transcription inhibition

Cells were cultured in YEP medium (1 % yeast extract, 2 % peptone, 40 μg ml^-1^ adenine) supplemented with 2 % glucose, or 2 % galactose, as stated, and at 30°C if not stated otherwise. For synchronization in G1-phase, 3 μg ml^-1^ α-factor mating pheromone (Merck, custom peptide WHWLQLKPGQPMY) was added every hour (a total of three additions) to cells growing logarithmically. For G2/M cell cycle arrest, benomyl (Merck, 381586) was added to the YEPD media for a final concentration 80 µg ml^-1^. A complete G2/M-arrest of logarithmically growing cells was achieved after 90 minutes at 30°C, or 120 minutes at 23°C. For G2/M-arrest following synchronization in G1-phase, cells were released into benomyl-containing YEPD for 60 minutes. For depletion of Scc2, Wpl1, Top1, Smc6, Nse4, Nse5 and Top2 in Figures 1A, 1D, 4A, S10C, auxin (3-indoleacetic acid, 1 mM final concentration, Merck, I2886) and doxycycline (5 μg ml^-1^ final concentration, Merck, D9891) were added to simultaneously degrade the proteins and inhibit their transcription, respectively. For degradation of Smc5 and Smc6 in Figure S9C, auxin (3-indoleacetic acid, Merck, I2886) was added at a final concentration of 1 mM. For transcription inhibition, thiolutin (Abcam, ab143556) was added to cell cultures for the final concentration of 20 µg ml^-1^ for 30 minutes. Cell cycle progression and arrests were confirmed using standard protocol for FACS analysis of ethanol-fixed, propidium iodide-stained cells.

### Protein extraction and Western blot

Depletion of HA-tagged Scc2, Wpl1, Top1, Smc6, Nse4, Nse5 and Top2 or degradation of MYC-tagged Smc5 and Smc6 were monitored by Western blot using anti-HA antibody clone 12CA5 (Roche, 1666606), or anti-c-Myc antibody (Merck, M4439), respectively, after protein extraction by trichloroacetic acid (TCA)-precipitation. Uncropped Western blot images are found in Figure S11.

### Chromatin immunoprecipitation, qPCR and ChIP-seq library preparation

Chromatin immunoprecipitation was performed as detailed in (Jeppsson *et al*., 2022). Briefly, *S. cerevisiae* cells were crosslinked with 1 % formaldehyde for 30 minutes at room temperature, followed by incubation at 4°C overnight. Chromatin was then sheared to a size of 300-500 bp by sonication (Bandelin Sonopuls HD 2070.2) and IP reactions containing anti-FLAG antibody (Merck, F1804) conjugated to Dynabeads Protein A (Invitrogen, 10002D), were allowed to proceed overnight at 4°C. After completing the immunoprecipitation and reversing crosslinks, the DNA was purified. ChIP-qPCR was performed using Fast SYBR Green (Applied Biosystems, 4385612) and primers listed in Table S2, using an Applied Biosystem 7500 Real-Time PCR System according to the manufacturer’s instructions. For ChIP-seq, DNA from ChIP and input fractions were prepared for sequencing using the NEBNext Ultra II DNA Library Prep Kit for Illumina (NEB, E7645). The libraries were sequenced using the HiSeq 2500 platform to generate single-end 65 bp reads. Sequenced reads were mapped to the *S. cerevisiae* genome using Bowtie2 version 2.4.1 with the default parameter set (Langmead and Salzberg, 2012). The numbers of total and mapped reads in each sample are listed in Table S3.

### ChIP-seq data analysis

To call chromosome arm peaks for Smc6 in *top2-4,* and Scc1 in wild-type cells, we identified bins in which the fold enrichment (ChIP / input) was more than 2.0 in the dataset 2017_036B_1367-X, and 2017_034B_234-X, respectively. Peaks overlapping with long terminal repeats, and pericentromeric regions (25 kb spanning each centromere), were excluded. The Smc5/6 binding sites in *top2-4* cells were used for the average enrichment-, and *cis* interaction analyses in Figures 1F, 2F, S3E, 4C, 7H, S10E. The cohesin binding sites in wild-type cells were used for the pile-up plot-and domain number analyses in Figures 7F, 7J, 7M, S10A. The 2691 ORFs used for the analysis of average Rpo21-FLAG enrichment in Figure 2B were selected to have average Rpo21 ChIP / input fold enrichment higher than 1.0 in wild-type cells, prior to the addition of thiolutin, *i.e.,* dataset 2017_035A_1958-2_0. We used DROMPAplus version 1.8. for normalizing, peak-calling, average enrichment analysis, and visualizing ChIP-seq data (Nakato and Sakata, 2020).

### RNA extraction, RT-qPCR and sequencing

Total RNA was prepared from 2.5×10^8^ yeast cells with Trizol (Invitrogen) and RNA extraction was performed according to the manufacture’s description. Briefly, 1 ml of Trizol reagent was added to the cells and cell disruption was achieved through vortexing (2 minutes) in the presence of glass beads. Then, 0.2 ml of chloroform was added to the cell suspension and incubated for 2 minutes at room temperature. Samples were centrifuged at 12,000 g for 15 minutes at 4°C and the aqueous phase was transferred to a fresh tube and mixed with 0.5 ml isopropanol. For RNA precipitation, samples were incubated at room temperature for 10 minutes and centrifuged at 12,000 g for 10 minutes at 4°C. RNA pellets were then washed with 70 % ethanol, air-dried and dissolved in 50 µl RNase-free water. cDNA for RT-qPCR was prepared using the High-Capacity cDNA Reverse Transcription Kit (ABI) according to the manufacture’s guidelines. qPCR for *MCR1*, *DBR1* and *ACT1* was performed using SYBR green (ABI) and primers listed in Table S2 on Applied Biosystem 7000 Real-Time PCR System, according to the manufacture’s guidelines. RNA-seq samples were prepared according to the manufacture’s standard protocol (TruSeq Standard Total RNA Sample Prep Kit, Illumina). Amplified cDNA was sequenced using the Illumina HiSeq platform (HiSeq2000) to generate single-end 50 bp reads. Sequence reads were aligned to the *S. cerevisiae* genome obtained from Saccharomyces Genome Database (http://www.yeastgenome.org/) using Bowtie (Langmead *et al*., 2009) with the default parameters. The numbers of total and mapped reads in each sample are listed in Table S3.

### Quantitative modeling of ChIP-seq data

Mathematical models to recapitulate Scc1 and Smc6 ChIP-seq profiles were constructed by a machine-learning algorithm. Since most of the Scc1 and Smc6 ChIP-seq peaks were located in the intergenic regions (IGRs) (Jeppsson *et al*., 2014), we focused on the Scc1 and Smc6 binding to IGRs. ChIP-seq fold-enrichment (FE) was calculated for each 10-bp bin in the genome, and the maximum FE within a 1 kb window centred on the midpoint of each IGR was used as the target variable. For predictor variables, 47 features associated with each IGR were calculated and used (summarized in Table S4). The data used for the feature calculation includes replication origin location in oriDB (Siow *et al*., 2012), replication fork merging zone (McGuffee *et al*., 2013), replication fork polarity (Clausen *et al*., 2015), as well as RNA-seq data from wild-type and *top2-4* cells arrested in G2/M after an S-phase at 35°C, restrictive temperature for the *top2-4* allele, obtained in this study. We built random forest regression models based on all or some of the features. The randomForest package in the R programming language was used for computing (Liaw and Wiener, 2002). The hyperparameter mtry was tuned by the tuneRF function. Model construction and evaluation were conducted by stratified fourfold cross-validation and iterated five times with differently split datasets. Spearman’s rank correlation coefficient (ρ) between the predicted and observed FE values was used for the evaluation of modelling accuracy.

### Smc5/6 complex purification and labelling

Hexameric (Smc5, Smc6, Nse1, Nse2, Nse3 and Nse4) and octameric (Smc5, Smc6, Nse1, Nse2, Nse3, Nse4, Nse5 and Nse6) *S. cerevisiae* Smc5/6 complexes were purified using the yeast overexpression protocol detailed in (Pradhan *et al*., 2023). Similarly, the fluorescently labelled Smc5/6 complexes which carry a C-terminal SNAP-tag on the Nse4 subunit were overexpressed, purified, and labelled as detailed in (Pradhan *et al*., 2023). Briefly, the Smc5/6 subunits under the control of Gal1-10 promoter were overexpressed in *S. cerevisiae* by adding 2 % galactose to exponentially growing yeast cells in YEP-lactate medium, and subsequently isolated by tandem affinity purification using IgG Sepharose 6 FF (VWR, 17-0969-01) and calmodulin Sepharose 4B (Merck). The eluate was concentrated using Vivaspin 20 100K MWCO ultrafiltration unit (Sartorius, VS2041), with simultaneous exchange to the buffer used for storage. For fluorescent labeling of the complexes, the eluate obtained from IgG Sepharose 6 FF was incubated with SNAP-Surface-Alexa Flour 647 (New England Biolabs, S9136S) at 4°C overnight. The resulting mixture was further purified and concentrated using Vivaspin20 ultrafiltration unit concomitant with buffer exchange.

### Single-molecule loop extrusion assay

The single-molecule experiment with supercoiled DNA was performed by building on methods described in (Pradhan *et al*., 2023) and (Kim *et al*., 2022). In essence, the process is split into three stages: surface functionalization, flow cell creation, and the loop extrusion assay with DNA supercoils. We used a custom-made HiLO (highly inclined optical light sheet) microscope for imaging, as per (Pradhan *et al*., 2023). During surface functionalization, we first prepared glass coverslips through silanization using a solution of 1% 3-[(2-aminoethyl)aminopropyl] trimethoxysilane in methanol and 5% glacial acetic acid. This was done after cleaning the coverslips with potassium hydroxide and acid piranha. Afterwards, we treated the silanized coverslips with a solution of 100mg/mL methoxyPEG-N-hydroxysuccinimide and 1 mg/mL biotin-PEG-N-hydroxysuccinimide in ice-cold borate buffer at pH 8.5. After overnight incubation, we rinsed the coverslips with milliQ water and dried them using nitrogen gas. This PEGylation was repeated five times before we sealed and stored the slides at −20 °C until use. For flow cell assembly, we drilled holes into slides to accommodate pipette tips. The same cleaning and functionalization processes were applied to these slides as to the coverslips. Flow channels were constructed using double-sided tape to sandwich the coverslip and slide together. One end of the resulting channel was connected to a syringe pump via a tube, and any openings, barring the drilled holes, were sealed to prevent sample leakage.

For the loop extrusion assay, we used 42 kb of coilable DNA (without nicks), with biotin ligated to both ends. We anchored the DNA to the coverslip by incubating 1 µM streptavidin in T50 buffer (40 mM Tris pH 7.5, 50 mM NaCl, 0.1 mM EDTA) within the channel, followed by a thorough wash. We then introduced a buffer at 3 µL/min containing the biotinylated DNA into the channel until we had a dense (but isolated, 100 DNA per 0.01 mm^2^) double-tethered DNA fill on the surface. We used Sytox Orange (SxO) for visualization and inducing twists in the DNA, introducing different supercoiling based on the presence of SxO in the imaging buffer. To find a SxO concentration inducing equivalent left-handed and right-handed twists, we plotted a titration curve (Figure S8A) using images of tethered DNA recorded at varying SxO concentrations using a 561 nm laser. We determined C1/2 ∼ 300 nM as the concentration where half of the intercalating sites are occupied, corresponding to half the maximum intensity. For negative supercoiling, DNA flowed at 800 nM SxO at 1.5 µL/min in the imaging buffer, occupying most intercalating sites. We maintained a final concentration of 300 nM SxO for both positive and negative supercoils during the loop extrusion assay.

Videos consisting of 10,000 frames at an acquisition speed of 100 ms per frame were recorded in the presence of 0.5 nM Smc5/6 hexamer combined with 2 mM ATP in the imaging buffer. Similarly, 1 nM hexamer tagged with Alexa647 and 2 mM ATP in the imaging buffer was introduced while alternating excitation between 561 nm and 640 nm lasers. For side-flow visualization, we employed a three ways channel, applying a flow of 20 µL/min perpendicular to the axis of the anchored DNA.

### Hi-C library preparation

Hi-C was performed as detailed in (Jeppsson *et al*., 2022). Briefly, *S. cerevisiae* cells were crosslinked with 3 % formaldehyde for 20 minutes at room temperature. Following spheroplasting and gentle lysis, restriction digestion of crosslinked chromatin by DpnII (NEB, R0543M) was performed overnight at 37°C, after which the restriction enzyme was inactivated by incubation at 62°C for 20 minutes. The presence of intact and individual DNA masses throughout the spheroplasting, digestion and ligation steps were confirmed by DAPI (4′,6-diamidino-2-phenylindole)-staining and microscopy. Marking and repairing DNA ends, proximity ligation, crosslink reversal, DNA shearing, size selection, biotin pull-down, preparation for Illumina sequencing, final amplification (15 cycles) and purification were then performed as in (Jeppsson *et al*., 2022). The Hi-C libraries were sequenced on Illumina HiSeq series with 150-bp paired-end sequencing according to the manufacturer’s recommendations. The numbers of total and mapped read pairs for each sample are listed in Table S5.

### Hi-C data analysis

The Hi-C data were processed using Juicer with the default parameter set (Durand et al., 2016). The sequenced reads were mapped to the *S. cerevisiae* genome obtained from Saccharomyces Genome Database (SGD) (http://yeastgenome.org/). The uniquely mapped read pairs were randomly resampled and arranged in the same numbers within sample groups (see Table S5). Contact matrices used for further analysis were coverage (sqrt)–normalized at 1-and 2-kb resolution with Juicer. The matrices were visualized by Juicebox (Robinson et al., 2018). Intrachromosomal contact frequency distribution was calculated using nonduplicated valid Hi-C contact pairs at genomic distances increasing by 1 kb.

Pile-up plots of pixels corresponding to pairs of specific sites in the contact matrices were calculated and normalized to the expected signal of global *cis* interactions at 1-kb resolution. Briefly, .cool format files were obtained from .hic format files by using hic2cool version 0.8.3 and the obtained matrices were normalized using the balance algorithm in cooler version 0.8.11 (Abdennur and Mirny, 2020). Expected Hi-C signals for *cis* interactions were calculated from these matrices using the expected-cis algorithm in cooltools version 0.5.4 (Open2C *et al*., preprint on bioRxiv: https://doi.org/10.1101/2022.10.31.514564). Pile-up plots were then calculated using coolpup.py with “--flank 15000 --mindist 0” option and plotted using plotpup.py in coolpup.py version 1.0.0 (Flyamer et al., 2020).

Domains were identified by using the Arrowhead algorithm in Juicer with coverage (sqrt) normalization at 1-and 2-kb resolution with “-m 300 -k VC_SQRT -r 2000” and “-m 200 -k VC_SQRT -r 1000” option. The Arrowhead detects the corners of the domains to identify their boundaries. The candidate domains at 1-and 2-kb resolution were merged, and the domains overlapping with Scc1 binding sites at both up-and downstream boundaries were used in the subsequent analysis.

The number of uniquely mapped *cis* and *trans* read pairs was normalized by total read number (read pair per kilobase). For *cis* interactions at Smc5/6 binding sites, uniquely mapped *cis* read pairs overlapping with these regions at either or both up-and downstream sites were obtained, and the number was normalized by total read number (read pair per million mapped read pairs).

## QUANTIFICATION AND STATISTICAL ANALYSIS

### ChIP-qPCR and Hi-C data

For ChIP-qPCR, two-sided *t*-test was used for analysis, and mean values from biological triplicates with error bars representing standard deviation are shown in the presented graphs. Binomial tests were used for the analysis of chromosomal *cis* / *trans* interactions, detected by Hi-C.

### Single-molecule loop extrusion assay

Microscopy images containing 10s of tethered DNAs in a field of view were examined to identify nicked and plectonemic DNA. DNAs exhibiting uniform intensity along their axis were categorized as nicked, whereas those with dynamic puncta were deemed plectonemic. Regions containing nicked and plectonemic DNA were isolated and saved as TIFF file for further quantification, using the Python based custom software described in (Pradhan *et al*., 2022). Specifically, we created kymographs by accumulating intensities on 11 pixels across the DNA axis and stacking each line of intensity for each image frame (see Figure 6B for an example). We located peak intensities corresponding to each punctum along the DNA axis using the “find_peaks” algorithm in Scipy (Virtanen *et al*., 2020). We calculated the area under each peak by summing intensities on 9 pixels around the peak to derive the puncta intensity (I_puncta_), while summing the remaining pixels gave the intensities outside the puncta (I_out_). We estimated puncta size as I_puncta_ (kb) = 42kb * I_puncta_/(I_puncta_+I_out_). We identified any punctum that localized to a single spot and increased in size upon Smc5/6 addition as a loop, determining its size using a similar method: I_loop_(kb) = 42kb * I_loop_/(I_loop_+I_out_). The loop size over time (t) provided the loop extrusion kinetics, and fitting the initial linear slope with the formula I_loop_ = k*t yielded the loop extrusion rate (k).

For labeled Smc5/6, we obtained kymographs for both the 561 nm (DNA) and 640 nm excitations. We extracted the intensities at the same positions as the DNA puncta or loops from the 640 nm excited kymograph, yielding time traces (see Figure 6F-G for an example) that facilitated estimating the number of hexamers involved in loop extrusion.

