## Supplemental Information for "Loop-extruding Smc5/6 organizes transcription-induced positive DNA supercoils"

**This PDF file includes:**

Table S1, S2, S3, S4, S5

Legends for Supplemental Figures S1 to S11

Supplemental Figures S1 to S11

Legends for Supplemental Movies S1 to S3

**Other Supplemental Information for this manuscript include the following:**

Movies S1, S2, S3

**Table S1. Yeast strains used in this study**

All strains are haploid and of W303 origin, carrying the genotype *ade2-1 trp1-1 can1-100 leu2-3,112 his3-11,15 ura3-1 RAD5*, with the modifications listed below.

| | |
|---|---|
| CB3067 | *MATa SMC6-6xHIS-3xFLAG-KAN ura3-1::ADH1-OsTIR1-9MYC::URA3,* |
| CB3541 | *MATa SMC6-6xHIS-3xFLAG-KAN Nat-tTA-tetO7-3HA-AID*-SCC2 ura3-1::ADH1-OsTIR1-9MYC::URA3 leu2-3,112::tetR'-SSN6::LEU2* |
| CB3594 | *MATa Nat-tTA-PtetO7-3HA-AID*-RAD61 SMC6-6xHIS-3xFLAG-KAN ura3-1::ADH1-OsTIR1-9MYC::URA3 leu2-3,112::tetR'-SSN6::LEU2* |
| CB3069 | *MATa SMC6-6xHIS-3xFLAG-KAN top2-4 ura3-1::ADH1-OsTIR1-9MYC::URA3 leu2-3,112::tetR'-SSN6::LEU2* |
| CB3543 | *MATa top2-4 SMC6-6xHIS-3xFLAG-KAN Nat-tTA-tetO7-3HA-AID*-SCC2 ura3-1::ADH1-OsTIR1-9MYC::URA3 leu2-3,112::tetR'-SSN6::LEU2* |
| CB3595 | *MATa Nat-tTA-PtetO7-3HA-AID*-RAD61 SMC6-6xHIS-3xFLAG-KAN top2-4 ura3-1::ADH1-OsTIR1-9MYC::URA3 leu2-3,112::tetR'-SSN6::LEU2* |
| CB1958 | *MATa RPO21-6HIS-3xFLAG:KAN* |
| CB1959 | *MATa RPO21-6HIS-3xFLAG:KAN top2-4* |
| CB173 | *MATa SMC6-6HIS-3xFLAG:KAN* |
| CB1367 | *MATa top2-4 SMC6-6HIS-3xFLAG:KAN* |
| CB234 | *MATa SCC1-6HIS-3xFLAG:KAN* |
| CB383 | *MATa top2-4 SCC1-6HIS-3xFLAG:KAN* |
| CB3300 | *MATa rpb1-1 SMC6-6xHIS-3xFLAG-KAN ura3-1::ADH1-OsTIR1-9MYC::URA3 leu2-3,112::tetR'-SSN6::LEU2* |
| CB3330 | *MATa Nat-tTA-tetO7-3HA-AID*-TOP2 ura3-1::ADH1-OsTIR1-9MYC::URA3 leu2-3,112::tetR'-SSN6::LEU2 SMC6-6HIS-3xFLAG:KAN* |
| CB3337 | *MATa rpb1-1 SMC6-6xHIS-3xFLAG-KAN Nat-tTA-tetO7-3HA-AID*-TOP2 ura3-1::ADH1-OsTIR1-9MYC::URA3 leu2-3,112::tetR'-SSN6::LEU2* |
| CB67 | *MATa* |
| CB1735 | *MATa pGPD-Dbr1::NAT pADH-Mcr1::KAN* |
| CB1764 | *MATa pGPD-Dbr1::NAT pADH-Mcr1::KAN SMC6-6HIS-3xFLAG:KAN* |
| CB617 | *MATa scc1-73 SMC6-6HIS-3xFLAG:KAN* |
| CB2790 | *MATa scc1-73 pGPD-Dbr1::NAT pADH-Mcr1::KAN SMC6-6HIS-3xFLAG:KAN* |

| | |
|---|---|
| CB2857 | *MATa pGPD-Dbr1::NAT pGAL1/10-Mcr1::TRP1* |
| CB2860 | *MATa SMC6-6HIS-3xFLAG:KAN pGPD-Dbr1::NAT pGAL1/10-Mcr1::TRP1* |
| CB3318 | *MATa HIS-tTA-tetO7-3HA-AID*-TOP1 ura3-1::ADH1-OsTIR1-9MYC::URA3 leu2-3,112::tetR'-SSN6::LEU2 SMC6-6HIS-3xFLAG:KAN* |
| CB3341 | *MATa SMC6-6xHIS-3xFLAG-KAN top2-4 HIS-tTA-tetO7-3HA-AID*-TOP1 ura3-1::ADH1-OsTIR1-9MYC::URA3 leu2-3,112::tetR'-SSN6::LEU2* |
| CB3573 | *MATa pep4del::HPH top1del::NAT leu2::LEU2pRS305-NSE1-pGAL1-10-NSE2 his3::HIS3pRS303-NSE3-pGAL1-10-NSE4 ura3::URA3pRS306-SMC5-pGAL1-10-SMC6-TAP* |
| CB4028 | *MATa pep4del::HPH top1del::NAT leu2::LEU2pRS305-NSE1-pGAL1-10-NSE2 his3::HIS3pRS303-NSE3-pGAL1-10-NSE4-6xHis-SNAP-KAN ura3::URA3pRS306-SMC5-pGAL1-10-SMC6-TAP* |
| CB3627 | *MATa ura3::pADH1-OsTIR1.9xMYC-URA3* |
| CB3859 | *MATa ura3::pADH1-OsTIR1-9xMyc-URA3 SMC5-AID*-9xMyc-NAT SMC6-AID*-9xMyc-NAT* |
| CB980 | *MATa top2-4* |
| CB3060 | *MATa ura3-1::ADH1-OsTIR1-9MYC::URA3 leu2-3, 112::tetR'-SSN6::LEU2* |
| CB3331 | *MATa Nat-tTA-tetO7-3HA-AID*-TOP2 ura3-1::ADH1-OsTIR1-9MYC::URA3 leu2-3,112::tetR'-SSN6::LEU2* |
| CB3412 | *MATa HIS-tetO7-SMC6-HA-IAA17-KAN HIS-tTA-tetO7-3HA-AID*-NSE5 HIS3-tTA-pTetO7-NSE4-6XHA-IAA17-KANMX ura3-1::ADH1-OsTIR1-9MYC::URA3 leu2-3,112::tetR'-SSN6::LEU2* |
| CB3612 | *MATa Nat-tTA-tetO7-3HA-AID*-TOP2 HIS-tetO7-SMC6-HA-IAA17-KAN HIS-tTA-tetO7-3HA-AID*-NSE5 HIS3-tTA-pTetO7-NSE4-6XHA-IAA17-KANMX ura3-1::ADH1-OsTIR1-9MYC::URA3 leu2-3, 112::tetR'-SSN6::LEU2* |

**Table S2. Primers used in this study**

| Purpose | Sequence |
|---|---|
| ChIP-qPCR at *CEN9* | CACGAATACGAGATACAGGGTAATGAA |
| | ACAGCTGAAGCTTGCCTCTGTATG |
| ChIP-qPCR at *HRB1-PET8* | AGACAAGAGGCCTCAGAAGGCTTA |
| | AACGGCTGGAGAAATGAGAGCGTA |
| ChIP-qPCR at *CYR1-SYS1* | ATCTTCAATGTCATCACGTACT |
| | AAATGGCAAGAACCTTATCTCT |
| ChIP-qPCR at *YGR127W-UTP8* | CATTCCGGTATCTCTTGTTGGA |
| | GCGTCTACTGTGGTTACTGATAA |
| ChIP-qPCR at *PMS1-SWS2* | ATACAGGGAGTGACAACACAAA |
| | CCACGTTCATATTCTTAATGGCTAAG |
| ChIP-qPCR at *UBP10-MRPL19* | CCTTCCATCTCCAATAAACTATGCC |
| | GACCAACCCGTGCTTTAGGAGAGTTA |
| ChIP-qPCR at *GCN1-HOS2* | GCTCCCAGCATTGTAGCTAAT |
| | ATGATGTGGTGGCAAGTCTC |
| ChIP-qPCR at *BPH1* ORF | CGGAAGTGGGTAAGTGCTATT |
| | ACTACCGGATTCTGTCATGTTT |
| ChIP-qPCR at *HED1-DAD1* | GCGAGGTATAAATCTGCGTCGCTA |
| | CAAGAATTTAACCTGGGCGGCT |
| ChIP-qPCR at *MCR1-DBR1* | GCATGGAGGAAAGTAATTGA |
| | AGCCAAGCCTTCTTGAGAGA |
| ChIP-qPCR at *MCR1* ORF | GCAAACCGTAACCAACATTCC |
| | TCGTGGGATTCCTCCTCTATT |
| ChIP-qPCR at *DBR1* ORF | GAGGCCGCAAATCTCTTAGTAG |
| | GGCCCAATGGAGTCGTTTAT |
| ChIP-qPCR at *ADH1* ORF | CTTCGTGACCACCGACTAATG |
| | GCCAAAGGCCAACGAATTG |
| ChIP-qPCR at *TDH3* ORF | TTGGGACTCTGAAAGCCATAC |
| | GTAACATCATCCCATCCTCCAC |
| ChIP-qPCR at *SPT16-CHC1* | ACCAGCAAGACCTTGATCTG |
| | GTCCTATGACCTTGACCGTTAG |

| | |
|---|---|
| ChIP-qPCR at *MPT5-YGL176C* | TTCCTGTCTTGGTTGGCTTTA |
| | CTTCTTGGTGTACAATTCCATTCTG |
| ChIP-qPCR at *GFD2-GRX1* | TCACAAGCACTCTTCCGACACACT |
| | AGGGAGACTGGTGAATTGGAGGAA |
| RT-qPCR of *MCR1* mRNA | AGCATGTTCCAGGTCCAAAG |
| | GCCCAAATTGTTCAAGATACCG |
| RT-qPCR of *DBR1* mRNA | TGATTCCAAATACGGCGAGAC |
| | GGTGATTCTCGTCCTTCCAC |
| RT-qPCR of *ACT1* mRNA | TCGAACAAGAAATGCAAACCG |
| | GGCAGATTCCAAACCCAAAAC |

### Table S3. ChIP- and RNA-sequencing statistics

**ChIP-seq statistics**

| | | | bowtie2 statistics | | | | | parse2wig statistics | |
|---|---|---|---|---|---|---|---|---|---|
| **ID** | **Description** | **Input sample** | **Sequenced reads** | **Mapped reads** | **%** | **Unmapped reads** | **%** | **Read depth** | **Genome coverage** |
| 2019_030B_3067-1_IP | Smc6-FLAG, IP | 2019_030B_3067-1_WCE | 7 633 828 | 7 429 543 | 97,3 | 204 285 | 2,7 | 55,5 | 0,99 |
| 2019_030B_3067-1_WCE | Smc6-FLAG, Input | - | 4 627 220 | 4 529 018 | 97,9 | 98 202 | 2,1 | 51,7 | 0,99 |
| 2019_030B_3541-1_IP | Smc6-FLAG Scc2-AID, IP | 2019_030B_3067-1_WCE | 7 591 726 | 7 365 465 | 97,0 | 226 261 | 3,0 | 56,7 | 1 |
| 2019_030B_3594-1_IP | Smc6-FLAG Wpl1-AID, IP | 2019_030B_3067-1_WCE | 7 664 428 | 7 478 071 | 97,6 | 186 357 | 2,4 | 53,3 | 1 |
| 2019_030B_3067-2_IP | Smc6-FLAG, IP | 2019_030B_3067-2_WCE | 7 270 727 | 7 064 037 | 97,2 | 206 690 | 2,8 | 45,0 | 0,99 |
| 2019_030B_3067-2_WCE | Smc6-FLAG, Input | - | 4 369 875 | 4 267 740 | 97,7 | 102 135 | 2,3 | 49,6 | 0,99 |
| 2019_030B_3541-2_IP | Smc6-FLAG Scc2-AID, IP | 2019_030B_3067-2_WCE | 6 846 971 | 6 673 466 | 97,5 | 173 505 | 2,5 | 45,5 | 0,99 |
| 2019_030B_3594-2_IP | Smc6-FLAG Wpl1-AID, IP | 2019_030B_3067-2_WCE | 7 251 940 | 7 058 626 | 97,3 | 193 314 | 2,7 | 41,8 | 0,99 |
| 2019_030B_3069-2_IP | Smc6-FLAG *top2-4*, IP | 2019_030B_3067-2_WCE | 7 206 559 | 7 027 423 | 97,5 | 179 136 | 2,5 | 52,2 | 0,99 |
| 2019_030B_3543-2_IP | Smc6-FLAG *top2-4* Scc2-AID, IP | 2019_030B_3067-2_WCE | 7 097 211 | 6 919 040 | 97,5 | 178 171 | 2,5 | 45,4 | 0,99 |
| 2019_030B_3595-2_IP | Smc6-FLAG *top2-4* Wpl1-AID, IP | 2019_030B_3067-2_WCE | 7 358 070 | 7 176 381 | 97,5 | 181 689 | 2,5 | 53,4 | 0,99 |
| 2017_032B_173-2_0_IP | Smc6-FLAG, 0 min thiolutin, IP | 2017_032B_173-2_0_Input | 2 343 161 | 2 296 550 | 98,0 | 46 611 | 2,0 | 17,8 | 0,99 |
| 2017_032B_173-2_0_Input | Smc6-FLAG, 0 min thiolutin, input | - | 2 404 776 | 2 357 245 | 98,0 | 47 531 | 2,0 | 28,7 | 0,99 |
| 2017_032B_173-2_3+_IP | Smc6-FLAG, 3 min thiolutin, IP | 2017_032B_173-2_3+_Input | 2 258 791 | 2 205 006 | 97,6 | 53 785 | 2,4 | 16,8 | 0,99 |
| 2017_032B_173-2_3+_Input | Smc6-FLAG, 3 min thiolutin, input | - | 2 191 836 | 2 149 080 | 98,1 | 42 756 | 2,0 | 24,5 | 1 |
| 2017_032B_173-2_15+_IP | Smc6-FLAG, 15 min thiolutin, IP | 2017_032B_173-2_15+_Input | 2 437 763 | 2 388 204 | 98,0 | 49 559 | 2,0 | 15,7 | 0,99 |
| 2017_032B_173-2_15+_Input | Smc6-FLAG, 15 min thiolutin, input | - | 2 273 393 | 2 231 100 | 98,1 | 42 293 | 1,9 | 24,9 | 1 |
| 2017_032B_173-2_30+_IP | Smc6-FLAG, 30 min thiolutin, IP | 2017_032B_173-2_30+_Input | 2 426 937 | 2 375 224 | 97,9 | 51 713 | 2,1 | 17,3 | 0,99 |
| 2017_032B_173-2_30+_Input | Smc6-FLAG, 30 min thiolutin, input | - | 2 203 467 | 2 160 766 | 98,1 | 42 701 | 1,9 | 24,0 | 1 |
| 2017_032B_173-2_30-_IP | Smc6-FLAG, 30 min DMSO, IP | 2017_032B_173-2_30-_Input | 2 201 071 | 2 155 315 | 97,9 | 45 756 | 2,1 | 15,9 | 0,99 |
| 2017_032B_173-2_30-_Input | Smc6-FLAG, 30 min DMSO, input | - | 2 128 979 | 2 085 882 | 98,0 | 43 097 | 2,0 | 23,4 | 0,99 |
| 2017_032B_1367-2_0+_IP | Smc6-FLAG *top2-4*, 0 min thiolutin, IP | 2017_032B_1367-2_0+_Input | 2 269 977 | 2 226 672 | 98,1 | 43 305 | 1,9 | 14,7 | 0,99 |
| 2017_032B_1367-2_0+_Input | Smc6-FLAG *top2-4*, 0 min thiolutin, input | - | 2 202 473 | 2 159 297 | 98,0 | 43 176 | 2,0 | 23,9 | 1 |
| 2017_032B_1367-2_3+_IP | Smc6-FLAG *top2-4*, 3 min thiolutin, IP | 2017_032B_1367-2_3+_Input | 2 372 123 | 2 327 585 | 98,1 | 44 538 | 1,9 | 17,4 | 0,99 |
| 2017_032B_1367-2_3+_Input | Smc6-FLAG *top2-4*, 3 min thiolutin, input | - | 2 236 727 | 2 194 347 | 98,1 | 42 380 | 1,9 | 24,5 | 1 |
| 2017_032B_1367-2_15+_IP | Smc6-FLAG *top2-4*, 15 min thiolutin, IP | 2017_032B_1367-2_15+_Input | 2 167 185 | 2 122 650 | 98,0 | 44 535 | 2,1 | 14,0 | 0,99 |
| 2017_032B_1367-2_15+_Input | Smc6-FLAG *top2-4*, 15 min thiolutin, input | - | 2 275 659 | 2 234 643 | 98,2 | 41 016 | 1,8 | 24,6 | 1 |
| 2017_032B_1367-2_30+_IP | Smc6-FLAG *top2-4*, 30 min thiolutin, IP | 2017_032B_1367-2_30+_Input | 2 328 975 | 2 284 392 | 98,1 | 44 583 | 1,9 | 16,1 | 0,99 |
| 2017_032B_1367-2_30+_Input | Smc6-FLAG *top2-4*, 30 min thiolutin, input | - | 2 438 677 | 2 392 013 | 98,1 | 46 664 | 1,9 | 28,2 | 0,99 |
| 2017_032B_1367-2_30-_IP | Smc6-FLAG *top2-4*, 30 min DMSO, IP | 2017_032B_1367-2_30-_Input | 2 251 661 | 2 212 832 | 98,3 | 38 829 | 1,7 | 14,6 | 0,99 |
| 2017_032B_1367-2_30-_Input | Smc6-FLAG *top2-4*, 30 min DMSO, input | - | 2 402 073 | 2 347 078 | 97,7 | 54 995 | 2,3 | 27,8 | 0,99 |
| 2017_032B_234-2_0_IP | Scc1-FLAG, 0 min thiolutin, IP | 2017_032B_173-2_0_Input | 2 017 090 | 1 965 080 | 97,4 | 52 010 | 2,6 | 18,8 | 0,97 |
| 2017_034B_234-2_3+_IP | Scc1-FLAG, 3 min thiolutin, IP | 2017_032B_173-2_3+_Input | 3 085 012 | 2 999 742 | 97,2 | 85 270 | 2,8 | 31,3 | 0,98 |
| 2017_034B_234-2_15+_IP | Scc1-FLAG, 15 min thiolutin, IP | 2017_032B_173-2_15+_Input | 3 061 849 | 2 980 572 | 97,4 | 81 277 | 2,7 | 30,6 | 0,97 |
| 2017_034B_234-2_30+_IP | Scc1-FLAG, 30 min thiolutin, IP | 2017_032B_173-2_30+_Input | 3 149 653 | 3 064 618 | 97,3 | 85 035 | 2,7 | 31,6 | 0,97 |
| 2017_032B_234-2_30-_IP | Scc1-FLAG, 30 min DMSO, IP | 2017_032B_173-2_30-_Input | 2 006 396 | 1 955 624 | 97,5 | 50 772 | 2,5 | 18,6 | 0,97 |
| 2017_032B_383-2_0_IP | Scc1-FLAG *top2-4*, 0 min thiolutin, IP | 2017_032B_1367-2_0+_Input | 2 040 558 | 1 990 272 | 97,5 | 50 286 | 2,5 | 21,1 | 0,98 |
| 2017_032B_383-2_3+_IP | Scc1-FLAG *top2-4*, 3 min thiolutin, IP | 2017_032B_1367-2_3+_Input | 2 194 298 | 2 134 245 | 97,3 | 60 053 | 2,7 | 23,2 | 0,98 |
| 2017_032B_383-2_15+_IP | Scc1-FLAG *top2-4*, 15 min thiolutin, IP | 2017_032B_1367-2_15+_Input | 2 091 036 | 2 031 917 | 97,2 | 59 119 | 2,8 | 21,8 | 0,98 |
| 2017_032B_383-2_30+_IP | Scc1-FLAG *top2-4*, 30 min thiolutin, IP | 2017_032B_1367-2_30+_Input | 2 215 938 | 2 155 574 | 97,3 | 60 364 | 2,7 | 22,9 | 0,98 |
| 2017_032B_383-2_30-_IP | Scc1-FLAG *top2-4*, 30 min DMSO, IP | 2017_032B_1367-2_30-_Input | 2 192 301 | 2 141 393 | 97,7 | 50 908 | 2,3 | 22,4 | 0,98 |
| 2017_035A_1958-2_0_IP | Rpo21-FLAG, 0 min thiolutin, IP | 2017_032B_173-2_0_Input | 3 411 009 | 3 356 896 | 98,4 | 54 113 | 1,6 | 41,7 | 0,98 |
| 2017_034B_1958-2_3+_IP | Rpo21-FLAG, 3 min thiolutin, IP | 2017_032B_173-2_3+_Input | 3 067 626 | 3 019 842 | 98,4 | 47 784 | 1,6 | 35,9 | 0,99 |
| 2017_034B_1958-2_15+_IP | Rpo21-FLAG, 15 min thiolutin, IP | 2017_032B_173-2_15+_Input | 3 155 403 | 3 104 822 | 98,4 | 50 581 | 1,6 | 35,1 | 0,99 |
| 2017_035A_1958-2_30+_IP | Rpo21-FLAG, 30 min thiolutin, IP | 2017_032B_173-2_30+_Input | 3 181 793 | 3 122 223 | 98,1 | 59 570 | 1,9 | 35,2 | 0,99 |
| 2017_036B_1958-2_30-_IP | Rpo21-FLAG, 30 min DMSO, IP | 2017_032B_173-2_30-_Input | 4 293 866 | 4 220 998 | 98,3 | 72 868 | 1,7 | 53,0 | 0,98 |
| 2017_036B_1959-2_0_IP | Rpo21-FLAG *top2-4*, 0 min thiolutin, IP | 2017_032B_1367-2_0+_Input | 3 781 905 | 3 717 779 | 98,3 | 64 126 | 1,7 | 47,8 | 0,98 |
| 2017_036B_1959-2_3+_IP | Rpo21-FLAG *top2-4*, 3 min thiolutin, IP | 2017_032B_1367-2_3+_Input | 3 841 352 | 3 780 070 | 98,4 | 61 282 | 1,6 | 46,7 | 0,99 |
| 2017_036B_1959-2_15+_IP | Rpo21-FLAG *top2-4*, 15 min thiolutin, IP | 2017_032B_1367-2_15+_Input | 3 679 139 | 3 623 142 | 98,5 | 55 997 | 1,5 | 43,8 | 0,99 |
| 2017_036B_1959-2_30+_IP | Rpo21-FLAG *top2-4*, 30 min thiolutin, IP | 2017_032B_1367-2_30+_Input | 4 791 861 | 4 704 953 | 98,2 | 86 908 | 1,8 | 57,2 | 0,99 |
| 2017_036B_1959-2_30-_IP | Rpo21-FLAG *top2-4*, 30 min DMSO, IP | 2017_032B_1367-2_30-_Input | 5 477 109 | 5 391 837 | 98,4 | 85 272 | 1,6 | 68,8 | 0,99 |
| 2019_030B_3067-3_IP | Smc6-FLAG, IP | 2019_030B_3067-3_WCE | 4 033 774 | 3 934 191 | 97,5 | 99 583 | 2,5 | 29,3 | 1 |
| 2019_030B_3067-3_WCE | Smc6-FLAG, Input | - | 4 675 154 | 4 573 491 | 97,8 | 101 663 | 2,2 | 53,3 | 0,99 |
| 2019_030B_3318-3_IP | Smc6-FLAG Top1-AID, IP | 2019_030B_3067-3_WCE | 4 155 725 | 4 033 773 | 97,1 | 121 952 | 2,9 | 33,6 | 0,99 |
| 2019_030B_3069-3_IP | Smc6-FLAG *top2-4*, IP | 2019_030B_3067-3_WCE | 4 071 737 | 3 974 564 | 97,6 | 97 173 | 2,4 | 30,0 | 1 |
| 2019_030B_3341-3_IP | Smc6-FLAG Top1-AID *top2-4*, IP | 2019_030B_3067-3_WCE | 4 140 712 | 4 047 964 | 97,8 | 92 748 | 2,2 | 40,2 | 0,99 |
| 2017_035A_173-X_IP | Smc6-FLAG, IP | 2017_035A_173-X_Input | 2 962 345 | 2 881 517 | 97,3 | 80 828 | 2,7 | 23,5 | 0,99 |
| 2017_035A_173-X_Input | Smc6-FLAG, Input | - | 3 117 430 | 3 046 218 | 97,7 | 71 212 | 2,3 | 36,4 | 0,99 |
| 2017_036B_1367-X_IP | Smc6-FLAG *top2-4*, IP | 2017_035A_173-X_Input | 3 710 052 | 3 611 303 | 97,3 | 98 749 | 2,7 | 25,6 | 0,99 |
| 2017_034B_234-X_IP | Scc1-FLAG, IP | 2017_035A_173-X_Input | 2 973 336 | 2 906 299 | 97,8 | 67 037 | 2,3 | 30,4 | 0,98 |
| 2017_034B_383-X_IP | Scc1-FLAG *top2-4*, IP | 2017_035A_173-X_Input | 3 237 130 | 3 165 783 | 97,8 | 71 347 | 2,2 | 35,5 | 0,97 |

**RNA-seq statistics**

| | | bowtie2 statistics | | | | |
|---|---|---|---|---|---|---|
| **ID** | **Description** | **Sequenced reads** | **Mapped reads** | **%** | **Unmapped reads** | **%** |
| CB67 | WT | 49 398 157 | 47 230 510 | 95,6 | 2 167 647 | 4,4 |
| CB980 | *top2-4* | 45 020 259 | 42 935 503 | 95,4 | 2 084 756 | 4,6 |

### Table S4. Features used in multivariable analysis

| feature group | feature ID | feature name | description | Spearman's rho for FE value | | | |
|---|---|---|---|---|---|---|---|
| | | | | Smc6-FLAG in top2-4 | Smc6-FLAG in WT | Scc1-FLAG in top2-4 | Scc1-FLAG in WT |
| A | a1 | flag.conv | Binary variable to indicate adjacent genes arranged in convergent orientation. | 0,328 | 0,287 | 0,482 | 0,497 |
| | a2 | flag.div | Binary variable to indicate adjacent genes arranged in divergent orientation. | -0,190 | -0,158 | -0,277 | -0,275 |
| | a3 | inward.lenSum.adj | Summed length of adjacent genes transcribed in an inward direction (bp). | 0,270 | 0,182 | 0,420 | 0,417 |
| | a4 | outward.lenSum.adj | Summed length of adjacent genes transcribed in an outward direction (bp). | -0,392 | -0,366 | -0,542 | -0,556 |
| | a5 | flag.conv.lenSum.adj | Summed length of adjacent genes, when they are in a convergent orientation (bp). | 0,345 | 0,295 | 0,505 | 0,520 |
| B | b1 | gene.len.longer | Length of adjacent, longer gene (bp). | -0,055 | -0,113 | -0,066 | -0,077 |
| | b2 | gene.len.shorter | Length of adjacent, shorter gene (bp). | -0,082 | -0,180 | -0,005 | -0,023 |
| | b3 | gene.len.sum | Summed length of adjacent genes (bp). | -0,070 | -0,148 | -0,051 | -0,066 |
| | b4 | gene.len.ratio | Ratio of longer adjacent gene to shorter adjoining one. | 0,057 | 0,111 | -0,040 | -0,029 |
| | b5 | rpkm.stronger | Transcription activity of adjacent, stronger gene (rpkm). | 0,089 | 0,196 | -0,052 | -0,042 |
| | b6 | rpkm.weaker | Transcription activity of adjacent, weaker gene (rpkm). | 0,009 | 0,004 | -0,007 | -0,004 |
| | b7 | rpkm.sum | Sum of transcription activities of adjacent genes (rpkm). | 0,085 | 0,184 | -0,049 | -0,039 |
| | b8 | rpkm.ratio | Ratio of transcription activity of stronger adjacent gene to weaker adjoining one. | 0,104 | 0,219 | -0,033 | -0,026 |
| C | c1 | inward.rpmSum.adj | Sum of rpm values (lengths of transcripts weighted by abundance) of adjacent genes transcribed in an inward direction. | 0,257 | 0,245 | 0,322 | 0,327 |
| | c2 | outward.rpmSum.adj | Sum of rpm values of adjacent genes transcribed in an outward direction. | -0,260 | -0,181 | -0,479 | -0,489 |
| D | d1 | inward.lenSum.2k | Summed length of (parts of) genes that are located within surrounding +/- 2-kb region and transcribed in an inward direction (bp). | 0,356 | 0,239 | 0,548 | 0,553 |
| | d2 | outward.lenSum.2k | Summed length of (parts of) genes that are located within surrounding +/- 2-kb region and transcribed in an outward direction (bp). | -0,417 | -0,407 | -0,570 | -0,588 |
| | d3 | inward.lenSum.5k | Summed length of (parts of) genes that are located within surrounding +/- 5-kb region and transcribed in an inward direction (bp). | 0,467 | 0,259 | 0,621 | 0,598 |
| | d4 | outward.lenSum.5k | Summed length of (parts of) genes that are located within surrounding +/- 5-kb region and transcribed in an outward direction (bp). | -0,490 | -0,384 | -0,647 | -0,638 |
| | d5 | inward.lenSum.10k | Summed length of (parts of) genes that are located within surrounding +/- 10-kb region and transcribed in an inward direction (bp). | 0,464 | 0,204 | 0,515 | 0,460 |
| | d6 | outward.lenSum.10k | Summed length of (parts of) genes that are located within surrounding +/- 10-kb region and transcribed in an outward direction (bp). | -0,461 | -0,304 | -0,545 | -0,510 |
| E | e1 | inward.rpmSum.2k | Summed rpm values of (parts of) genes that are located within surrounding +/- 2-kb region and transcribed in an inward direction (bp). | 0,275 | 0,271 | 0,331 | 0,350 |
| | e2 | outward.rpmSum.2k | Summed rpm values of (parts of) genes that are located within surrounding +/- 2-kb region and transcribed in an outward direction (bp). | -0,216 | -0,138 | -0,423 | -0,431 |
| | e3 | inward.rpmSum.5k | Summed rpm values of (parts of) genes that are located within surrounding +/- 5-kb region and transcribed in an inward direction (bp). | 0,284 | 0,247 | 0,296 | 0,299 |
| | e4 | outward.rpmSum.5k | Summed rpm values of (parts of) genes that are located within surrounding +/- 5-kb region and transcribed in an outward direction (bp). | -0,201 | -0,083 | -0,373 | -0,366 |
| | e5 | inward.rpmSum.10k | Summed rpm values of (parts of) genes that are located within surrounding +/- 10-kb region and transcribed in an inward direction (bp). | 0,252 | 0,201 | 0,206 | 0,198 |
| | e6 | outward.rpmSum.10k | Summed rpm values of (parts of) genes that are located within surrounding +/- 10-kb region and transcribed in an outward direction (bp). | -0,167 | -0,039 | -0,291 | -0,270 |
| F | f1 | ATrate.start2stop | AT-richness of corresponding intergenic region (%). | 0,308 | 0,208 | 0,383 | 0,391 |
| | f2 | ATskew.start2stop | AT skewness (absolute value) of corresponding intergenic region. | 0,016 | -0,003 | 0,028 | 0,028 |
| | f3 | GCskew.start2stop | GC skewness (absolute value) of corresponding intergenic region. | 0,033 | 0,021 | 0,062 | 0,061 |
| | f4 | igr.len | Length of corresponding intergenic region (bp). | -0,091 | 0,053 | -0,241 | -0,234 |
| | f5 | ATrate.1kb | AT-richness of 1-kb region centered on the midpoint of corresponding intergenic region (%). | 0,328 | 0,283 | 0,297 | 0,308 |
| | f6 | ATskew.1kb | AT skewness (absolute value) of 1-kb region centered on the midpoint of corresponding intergenic region. | -0,041 | -0,061 | -0,052 | -0,061 |
| | f7 | GCskew.1kb | GC skewness (absolute value) of 1-kb region centered on the midpoint of corresponding intergenic region. | -0,029 | -0,025 | -0,021 | -0,017 |
| G | g1 | dist.nearestOri.db | Distance to the nearest replication origin (confirmed origins in oriDB) (bp).* | 0,031 | -0,057 | 0,004 | -0,018 |
| | g2 | rep.dir.bias | Degree of directional bias in replication fork movement, based on HydEn-seq data.* | -0,167 | -0,100 | -0,095 | -0,081 |
| | g3 | dif.rep.dir | Derivative of g2. * | -0,027 | 0,038 | -0,018 | -0,001 |
| | g4 | dist.nearestFMZ | Distance to the nearest replication termination zone (FMZ, McGuffee et al.).* | -0,116 | 0,023 | -0,021 | 0,001 |
| | g5 | dist.nearestFMZ.TER | Distance to the nearest replication termination zone (TER-FMZ, McGuffee et al.).* | -0,101 | -0,077 | -0,116 | -0,128 |
| | g6 | collision2longerGene | Degree of replication-transcription collision for adjacent, longer gene.* | 0,059 | 0,012 | 0,073 | 0,066 |
| | g7 | collision2strongerGen | Degree of replication-transcription collision for adjacent, stronger gene.* | 0,030 | -0,001 | 0,049 | 0,048 |
| | g8 | len.collideGene | Legth of (longer) gene that is transcribed in an opposite direction to the majority of replication fork (bp). | 0,073 | -0,010 | 0,105 | 0,098 |
| | g9 | rpkm.collideGene | Rpkm value of (more active) gene that is transcribed in an opposite direction to the majority of replication fork. | 0,095 | 0,093 | 0,076 | 0,084 |
| H | h1 | chr.len | Length of the corresponding chromosome (bp). | 0,140 | 0,285 | -0,060 | -0,055 |
| | h2 | dist.nearestCen | Distance to centromere (bp).* | -0,080 | 0,010 | -0,159 | -0,166 |
| | h3 | dist.nearestTel | DIstance to the nearest telomere (bp).* | 0,278 | 0,264 | -0,023 | -0,023 |
| | h4 | dist.TelSameArm | Distance to the telomere of the same chromosome arm (bp).* | 0,304 | 0,255 | 0,018 | 0,020 |

* The reference point is the mid-point of the coresponding IGR.

### Table S5. Hi-C sequencing statistics

| ID | Description | Total number of read pairs | Number of uniquely mapped read pairs | Number of read pairs after filtering | Number of *cis* - read pairs (percentage) | Number of *trans* - read pairs (percentage) |
|---|---|---|---|---|---|---|
| KJ48 | WT | 93 147 908 | 35 761 005 | 27 559 838 | 23 103 550 (83,8%) | 4 456 288 (16,2%) |
| KJ49 | WT | 75 393 887 | 35 761 005 | 27 142 697 | 22 607 497 (83,3%) | 4 535 200 (16,7%) |
| KJ50 | Smc5-AID Smc6-AID | 107 038 329 | 35 761 005 | 26 100 783 | 20 296 597 (77,8%) | 5 804 186 (22,2%) |
| KJ51 | Smc5-AID Smc6-AID | 87 713 224 | 35 761 005 | 26 045 412 | 20 045 102 (77,0%) | 6 000 310 (23,0%) |
| KJ56 | WT | 113 957 255 | 35 761 005 | 27 110 271 | 20 370 085 (75,1%) | 6 740 186 (24,9%) |
| KJ57 | WT | 78 616 080 | 35 761 005 | 26 850 477 | 20 154 419 (75,1%) | 6 696 058 (24,9%) |
| KJ60 | *top2-4* | 50 933 723 | 35 761 005 | 26 886 024 | 21 547 838 (80,1%) | 5 338 186 (19,9%) |
| KJ61 | *top2-4* | 103 532 105 | 35 761 005 | 27 551 659 | 22 367 253 (81,2%) | 5 184 406 (18,8%) |
| KJ6 | WT | 130 461 549 | 66 000 000 | 53 966 058 | 37 077 612 (68,7%) | 16 887 479 (31,3%) |
| KJ7 | Top2-AID | 101 439 833 | 66 000 000 | 52 993 845 | 39 057 222 (73,7%) | 13 935 285 (26,3%) |
| KJ8 | Smc6-AID Nse4-AID Nse5-AID | 104 087 372 | 66 000 000 | 53 566 606 | 37 786 787 (70,5%) | 15 778 844 (29,5%) |
| KJ10 | Top2-AID Smc6-AID Nse4-AID Nse5-AID | 94 640 454 | 66 000 000 | 52 251 004 | 37 391 395 (71,6%) | 14 858 253 (28,4%) |

#### SUPPLEMENTAL FIGURE LEGENDS

**Supplemental Figure 1. Schematics of DNA supercoiling, and FACS analysis of Scc2- and Wpl1-depleted cells. Related to Figure 1.**

(A) Positive supercoiling accumulates ahead of, and negative supercoiling behind, transcribing RNA pol II. (B) Positive supercoils also accumulate ahead of the replication fork. Fork rotation can allow the channeling of positive supercoils ahead of the fork into sister chromatid intertwinings (SCIs) behind the fork. (C) Positively supercoiled DNA can take the form as overtwisted DNA, or positively supercoiled plectonemic DNA if the increased twist is converted to writhe. The positively supercoiled plectoneme is a left-handed helix. (D) FACS analysis displaying the G2/M-arrest of the wild-type, Scc2- or Wpl1-depleted cells used in the analysis displayed in panel E and in Figure 1A-C. (E) Averaged Smc6-FLAG enrichment in 25 kb regions spanning each centromere in wild-type, Scc2- or Wpl1-depleted G2/M-arrested cells, as determined by ChIP-seq analysis. (F) FACS analysis of the wild-type and *top2-4* mutant cells, with or without Scc2- or Wpl1-depletion, used in the analysis displayed in Figure 1D-G.

**Supplemental Figure 2. RNA pol II, but not cohesin, dissociates from chromosomes rapidly following thiolutin treatment. Related to Figure 2.**

(A) Rpo21-FLAG and (B) Scc1-FLAG enrichment along the arm of chromosome 7, 100 - 200 kb from the left telomere, in wild-type (upper five tracks) and *top2-4* mutated (lower five tracks) G2/M-arrested cells, with or without preceding thiolutin-treatment during indicated periods, as determined by ChIP-seq. Annotations as in Figure 1B.

**Supplemental Figure 3. Smc5/6, but not cohesin, dissociates rapidly from chromosomes following thiolutin treatment. Related to Figure 2.**

(A) FACS analysis of the wild-type and *top2-4* mutant cells, with or without preceding thiolutin-treatment during indicated periods, used in the analysis displayed in Figure 1C-G. (B-C) Averaged Smc6-FLAG enrichment in 25 kb regions spanning each centromere in (B) wild-type and (C) *top2-4* G2/M-arrested cells, with or without indicated periods of thiolutin-treatment. (D) Smc6-FLAG enrichment within the *BPH1* open reading frame, a Smc5/6-"non-binding site", and in the intergenic regions between the *GCN1-HOS2* convergent gene pair as determined by ChIP-qPCR in G2/M-arrested cells, with or without preceding thiolutin-treatment during indicated periods. *N=1*. (E) Averaged Smc6-FLAG enrichment at Smc5/6 binding sites, based on the analysis presented in Figure 2C and G. (F) Scc1-FLAG enrichment in the centromeric region of chromosome 10, 420 - 450 kb from the left telomere, in G2/M-arrested wild-type cells, with or without indicated periods of thiolutin-treatment. Annotations as in Figure 1B. (G) Scc1-FLAG enrichment within the *BPH1* open reading frame, a cohesin-"non-binding site", and in the intergenic regions between the *GCN1-HOS2* convergent gene pair as determined by ChIP-qPCR in G2/M-arrested cells, with or without preceding thiolutin-treatment during indicated periods. *N=1*. (H) Smc6-FLAG enrichment at centromere 9, in intergenic regions between indicated convergent gene pairs, and within the *BPH1* open reading frame, a Smc5/6-"non-binding site", as determined by ChIP-qPCR analysis of wild-type and Top2-depleted cells, with or without inactivation of RNA pol II function using the temperature sensitive *rpb1-1* allele. *N=3,* n.s.*: p>0.05*, ∗ *: p≤0.05,* ∗∗ *: p≤0.01.*

**Supplemental Figure 4. Overexpression of *MCR1* and *DBR1* does not affect cellular growth. Related to Figure 3.**

(A) Scc1-FLAG enrichment in wild-type cells (upper track), and Smc6-FLAG enrichment in wild-type or *top2-4* mutated cells (middle tracks) on chromosome 11, 100 - 200 kb from the left telomere. Note the low level of Smc6-FLAG enrichment in the *MCR1-DBR1* intergenic region (dotted line) in both cell types. Annotations as in Figure 1B, with an additional highlight of the *MCR1* and *DBR1* open reading frames in lilac and green, respectively. (B) Overexpression of *MCR1* and *DBR1* does not perturb cell cycle progression. FACS profiles of cells over-expressing *MCR1* and *DBR1*, or not, arrested in the G1-phase and released into the cell cycle at 35°C. (C). Tenfold serial dilutions of cells overexpressing *MCR1* and *DBR1*, or not, on solid medium containing 0.015 % methyl methane sulfonate (MMS), 100 mM hydroxyurea (HU) or 10 µg/ml benomyl that convey S-phase DNA damage, replication fork blockage, or microtubule destabilization, respectively. The plates were incubated for three (MMS and HU) or four (benomyl) days at 25°C. Note that both cell-types grow equally well. The *mms21-CH*, *rad50Δ* and *mad2Δ* strains were included as controls for cells with increased sensitivity to MMS, HU and benomyl, respectively.

**Supplemental Figure 5. Smc5/6, but not cohesin, is recruited site-specifically in between a highly expressed convergently oriented gene pair. Related to Figure 3.**

(A) ChIP-on-chip of Smc6-FLAG in cells with endogenous expression of *MCR1* and *DBR1* (left) or overexpression of the two genes (right). ChIP-on-chip was performed on cells arrested in G2/M by nocodazole at 35°C, after passage through a synchronous S-phase. Regions 100 - 200 kb and 400 - 500 kb from the left telomere of chromosome 11 are shown. The Y-axis shows fold enrichment of ChIP / whole cell extract in $log_2$ scale, while the X-axis displays chromosomal positions. Blue horizontal bars denote open reading frames and red vertical lines denote replication origins (ARS). (B) Scc1-FLAG enrichment at centromere 9, in the intergenic regions between the *MCR1-DBR1* convergent gene pair, and within *MCR1*, *DBR1*, *ADH1*, *TDH3* open reading frames as indicated. The data was obtained by ChIP-qPCR analysis of cells with or without *MCR1* and *DBR1* promoter replacement. *N=3*. (C) Relative expression level of *MCR1* and *DBR1* as compared to *ACT1*, with or without *GAL1-10* and *GPD* promoter replacement, respectively, as determined by RT-qPCR. (D) FACS analysis of $P_{Gal1\text{-}10}$-*ADH1* $P_{GPD}$-*MCR1* cells arrested in G1-phase in YEP-media containing 2% raffinose, with subsequent addition of glucose or galactose (final concentration 2%), used in the analysis displayed in Figure 3G. (E) Smc6-FLAG enrichment at centromere 9, at intergenic regions between convergent gene pairs, and within *MCR1* and *DBR1* open reading frames, as indicated. The data was obtained by ChIP-qPCR analysis of G2/M-arrested $P_{Gal1\text{-}10}$-*ADH1* $P_{GPD}$-*MCR1* cells, treated with glucose or galactose (final concentration 2%). *N=3*. (F) FACS analysis of $P_{Gal1\text{-}10}$-*ADH1* $P_{GPD}$-*MCR1* cells arrested in G2/M-phase in YEP-media containing 2% raffinose, with subsequent addition of glucose or galactose (final concentration 2%), used in the analysis displayed in panel E.

**Supplemental Figure 6. FACS analysis of G2/M-arrest of cells experiencing Top1 and Top2 inactivation, related to Figure 4.**

(A) FACS analysis displaying the G2/M-arrest of the wild-type, Top1-AID, *top2-4*, and the Top1-AID *top2-4* cells used in the analysis displayed in Figure 4B-D.

**Supplemental Figure 7. Prediction of cohesin chromosomal enrichment in wild-type and *top2-4* cells. Related to Figure 5.**

(A) Evaluation of the accuracy of prediction models for Scc1-FLAG enrichment in *top2-4* and wild-type cells using the indicated groups of features,m as in Figure 5A. Four-fold cross-validation was repeated five times with differently partitioned datasets, and the resulting correlation coefficients are plotted. The horizontal line indicates the mean, and whiskers show the 95% confidence interval. (B) Representative prediction results of analysis in (A). Observed and predicted FE values, $FE_{obs}$ and $FE_{pred}$, respectively, were plotted in $log_2$ scale.

**Supplemental Figure 8. Dimers of hexameric Smc5/6 loop extrudes on positively supercoiled DNA. Related to Figure 6.**

(A) The intensity of DNA with different SYTOX Orange concentrations. $C_{1/2}$ (300 nM), the concentration at which half of the maximum occupancy was bound by the intercalator is indicated, and was used for generation and visualization of positive and negative supercoils. (B-C) Kymographs of an example of (B) positive and a (C) negative supercoiled DNA, respectively. (D) Size and number of plectonemes following addition of hexameric Smc5/6 complex, corresponding to Figure 6A-C. (E) Violin plots showing Smc5/6 loop extrusion rates on nicked and positively supercoiled (+ve SC) DNA substrate. (F) The number of bleaching steps of labeled hexameric Smc5/6 bound at the initiation of loop extrusion on positively supercoiled DNA. The dashed bars correspond to theoretical values assuming that these complexes were dimers, with the labeling efficiency of 72 %. (G) An example of kymographs for loop extrusion by labeled hexameric Smc5/6 (bottom) on nicked DNA (top). (H) The time traces of the DNA loop size (top) and the intensity of labeled Smc5/6 (bottom) corresponding to the kymographs in panel (G). (I) Kymographs of an example of binding and translocation events using labeled octameric (Nse5/6-containing) Smc5/6 on positively supercoiled (top) and nicked (bottom) DNA. Yellow arrows indicate the binding events and white arrows indicate the end position of translocation events by Smc5/6 on DNA. (J) The average number of octameric Smc5/6 binding events on positively supercoiled versus nicked DNA per DNA molecule. (K) The fraction of octameric complexes that bound, translocated and ended up at DNA ends on nicked or positively supercoiled DNA.

**Supplemental Figure 9. Smc5/6 accumulates on, and promotes association of, supercoiled loci on entangled chromosomes. Related to Figure 7.**

(A) FACS analysis of wild-type Smc5-AID and Smc6-AID cells used in the analysis displayed in Figure 7A-F. (B) Western blot analysis showing the depletion of Smc5 and Smc6 related to panel A, and Figure 7A-F. (C) FACS analysis of wild-type and *top2-4* cells used in the analysis displayed in panel D and Figure 7G-J. (D) Normalized Hi-C contact maps (2 kb binning) showing *cis* interactions along the arm of chromosome 2, 450 - 540 kb from left telomere, in G2/M-arrested wild-type and *top2-4* cells after an S-phase at 35°C, restrictive temperature for the *top2-4* allele. Lines on top and to the left of the panels: green = cohesin binding sites in *top2-4*, blue = Smc5/6 binding sites in *top2-4*, light blue arrow: example of increased *cis* interactions between Smc5/6 binding sites in *top2-4*. (E-F) Smc6-FLAG (upper two tracks) and Scc1-FLAG (lower two tracks) enrichment in wild-type or *top2-4* mutated cells, on (E) chromosome 7, 80 - 160 kb from the left telomere, and (F) chromosome 2, 450 - 540 kb from the left telomere. These ChIP-seq maps correspond to the regions displayed in Figure 7I and L, and panel D and S10C, respectively. Annotations as in Figure 1B, with the addition that the blue arrows indicate the Smc5/6 sites between which an increase in *cis* interactions was detected in the absence of Top2 function.

**Supplemental Figure 10. Smc5/6-dependent *cis* interactions detected in Top2-AID cells. Related to Figure 7.**

(A) Pile-up plots of averaged *cis* interactions between all pairs of cohesin sites situated 20 - 40 kb apart on chromosome arms in wild-type and *top2-4* cells. (B) FACS analysis of wild-type, triple Smc6-AID Nse4-AID Nse5-AID, Top2-AID, and quadruple Top2-AID Smc6-AID Nse4-AID Nse5-AID cells used in the analysis displayed in Figure 7K-M. (C) Western blot analysis showing the depletion of Top2-AID Smc6-AID Nse4-AID Nse5-AID related to panel B, and Figure 7K-M. (D) Normalized Hi-C contact maps (2 kb binning) showing *cis* interactions along the arm of chromosome 2, 450 - 540 kb from left telomere, in wild-type, Smc6-AID Nse4-AID Nse5-AID triple depletion, Top2-AID, and Top2-AID Smc6-AID Nse4-AID Nse5-AID quadruple depletion cells. Lines on top and to the left of the panels: green = cohesin binding sites in *top2-4* mutant, blue = Smc5/6 binding sites in *top2-4* mutant, light blue arrow: example of increased Smc5/6-dependent *cis* interactions between Smc5/6 binding sites in Top2-AID cells. (E) Quantification of *cis* interactions (>10 kb) anchored at Smc5/6 binding sites on entangled chromosome arms in wild-type, Smc6-AID Nse4-AID Nse5-AID triple depletion, Top2-AID, and Top2-AID Smc6-AID Nse4-AID Nse5-AID quadruple depletion cells.

**Supplemental Figure 11. Uncropped Western blot images of protein depletion.**

(A-E) Ponceau and stain-free gel (left), and uncropped Western blot (right) images of protein depletion used in Figures 1A, 1D, 4A, S9B and S10C, respectively.

**Figure S1**

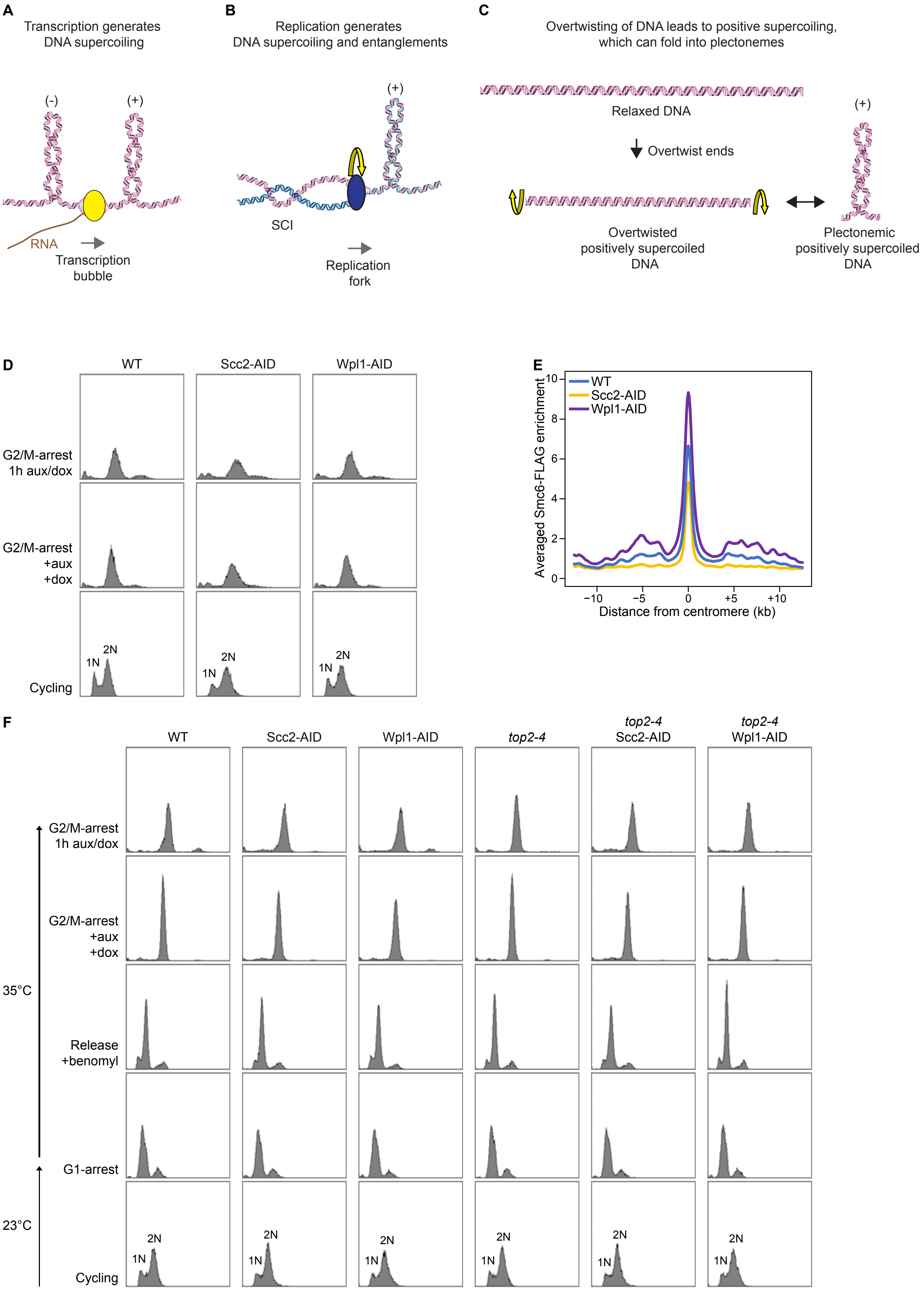
A
Transcription generates DNA supercoiling
(-)
(+)
RNA
Transcription bubble
B
Replication generates DNA supercoiling and entanglements
(+)
SCI
Replication fork
C
Overtwisting of DNA leads to positive supercoiling, which can fold into plectonemes
Relaxed DNA
Overtwist ends
(+)
Overtwisted positively supercoiled DNA
Plectonemic positively supercoiled DNA
D
WT
Scc2-AID
Wpl1-AID
G2/M-arrest 1h aux/dox
G2/M-arrest +aux +dox
Cycling
1N
2N
E
WT
Scc2-AID
Wpl1-AID
Averaged Smc6-FLAG enrichment
10
8
6
4
2
0
−10
−5
0
+5
+10
Distance from centromere (kb)
F
WT
Scc2-AID
Wpl1-AID
top2-4
top2-4 Scc2-AID
top2-4 Wpl1-AID
G2/M-arrest 1h aux/dox
G2/M-arrest +aux +dox
35°C
Release +benomyl
G1-arrest
23°C
Cycling
1N
2N

**Figure S2**

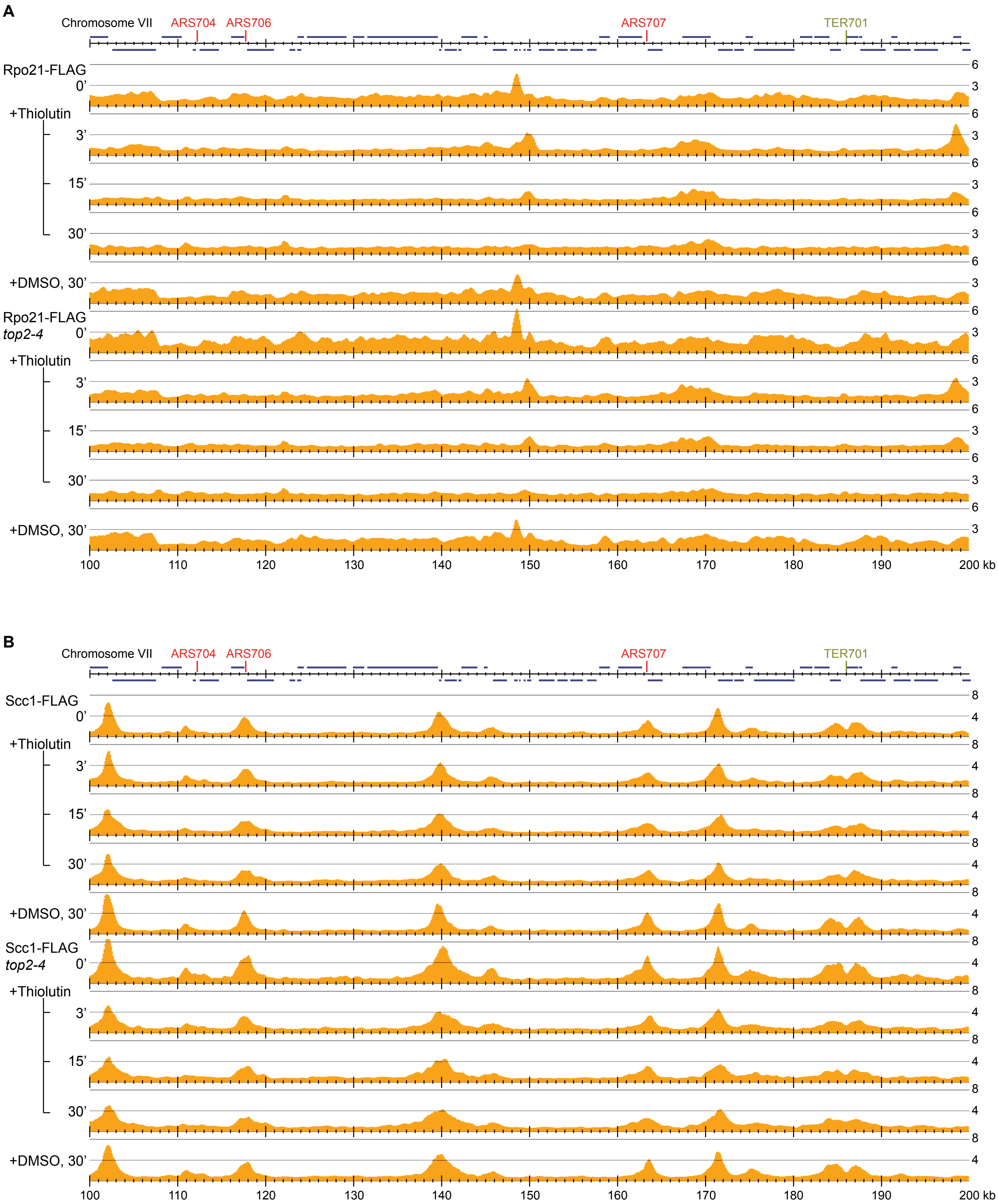
A
Chromosome VII
ARS704
ARS706
ARS707
TER701
Rpo21-FLAG
0'
+Thiolutin
3'
15'
30'
+DMSO, 30'
Rpo21-FLAG
top2-4
0'
+Thiolutin
3'
15'
30'
+DMSO, 30'
6
3
100
110
120
130
140
150
160
170
180
190
200 kb
B
Chromosome VII
ARS704
ARS706
ARS707
TER701
Scc1-FLAG
0'
+Thiolutin
3'
15'
30'
+DMSO, 30'
Scc1-FLAG
top2-4
0'
+Thiolutin
3'
15'
30'
+DMSO, 30'
8
4
100
110
120
130
140
150
160
170
180
190
200 kb

**Figure S3**

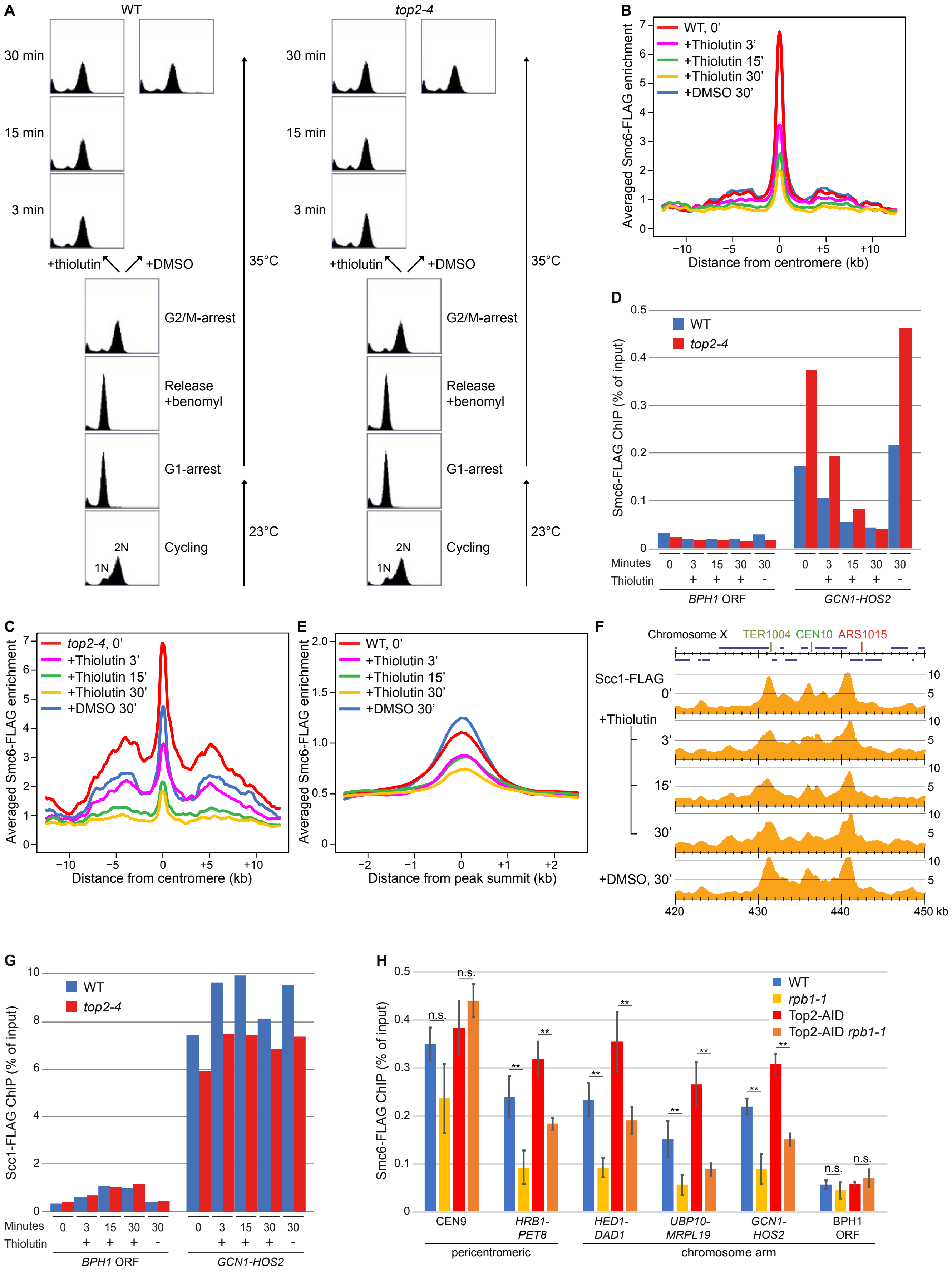

A
WT
top2-4
30 min
15 min
3 min
+thiolutin
+DMSO
35°C
23°C
G2/M-arrest
Release
+benomyl
G1-arrest
Cycling
2N
1N
B
Averaged Smc6-FLAG enrichment
WT, 0’
+Thiolutin 3’
+Thiolutin 15’
+Thiolutin 30’
+DMSO 30’
Distance from centromere (kb)
C
top2-4, 0’
D
Smc6-FLAG ChIP (% of input)
WT
top2-4
Minutes
Thiolutin
BPH1 ORF
GCN1-HOS2
E
Distance from peak summit (kb)
F
Chromosome X
TER1004
CEN10
ARS1015
Scc1-FLAG
0’
+Thiolutin
3’
15’
30’
+DMSO, 30’
420
430
440
450 kb
G
Scc1-FLAG ChIP (% of input)
H
rpb1-1
Top2-AID
Top2-AID rpb1-1
n.s.
**
CEN9
HRB1-PET8
HED1-DAD1
UBP10-MRPL19
GCN1-HOS2
BPH1 ORF
pericentromeric
chromosome arm

**Figure S4**

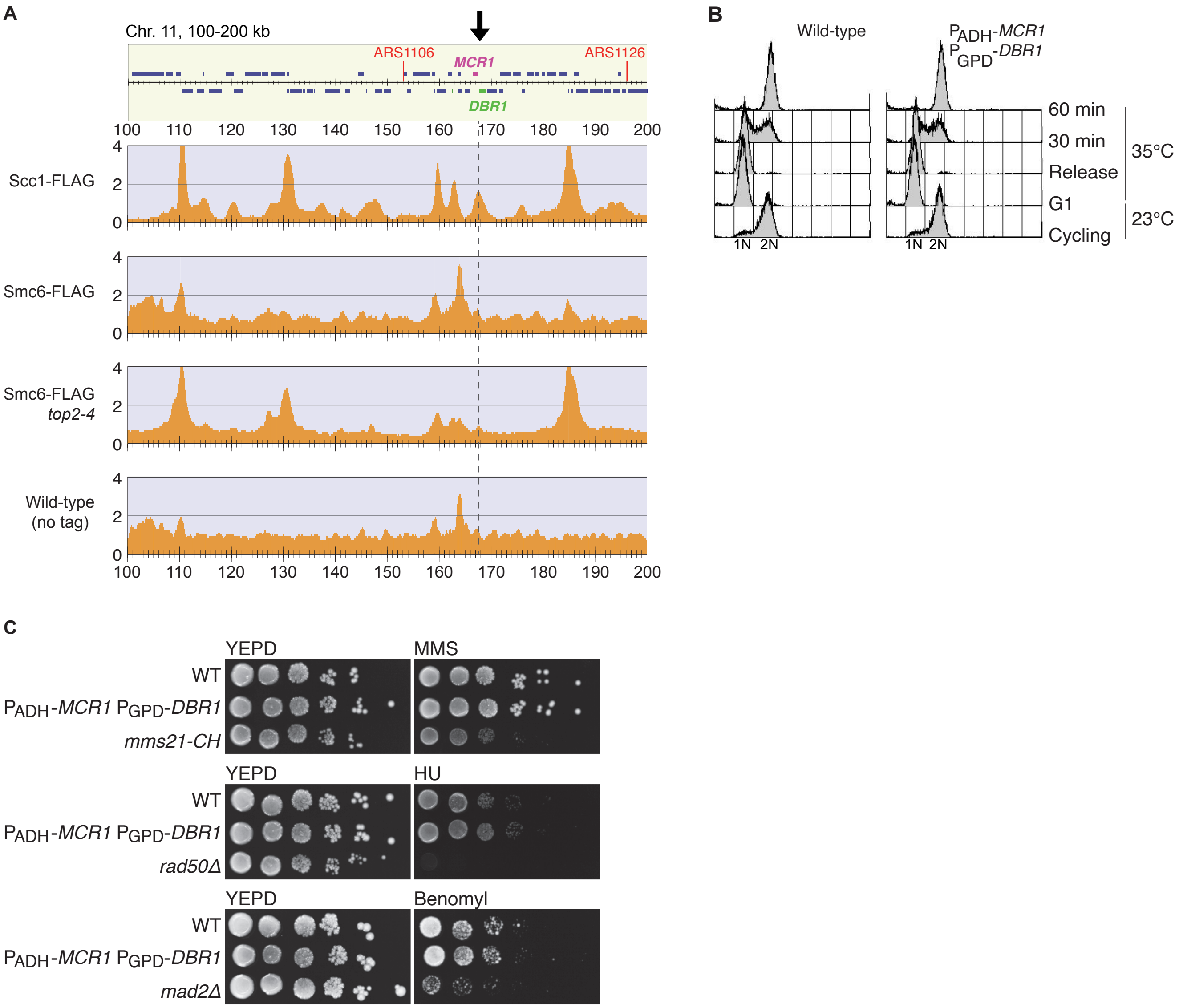
A
Chr. 11, 100-200 kb
ARS1106
MCR1
ARS1126
DBR1
100 110 120 130 140 150 160 170 180 190 200
Scc1-FLAG
Smc6-FLAG
Smc6-FLAG
top2-4
Wild-type
(no tag)
0
2
4
B
Wild-type
PADH-MCR1
PGPD-DBR1
60 min
30 min
Release
G1
Cycling
35°C
23°C
1N 2N
C
YEPD
MMS
HU
Benomyl
WT
PADH-MCR1 PGPD-DBR1
mms21-CH
rad50Δ
mad2Δ

#### Figure S5

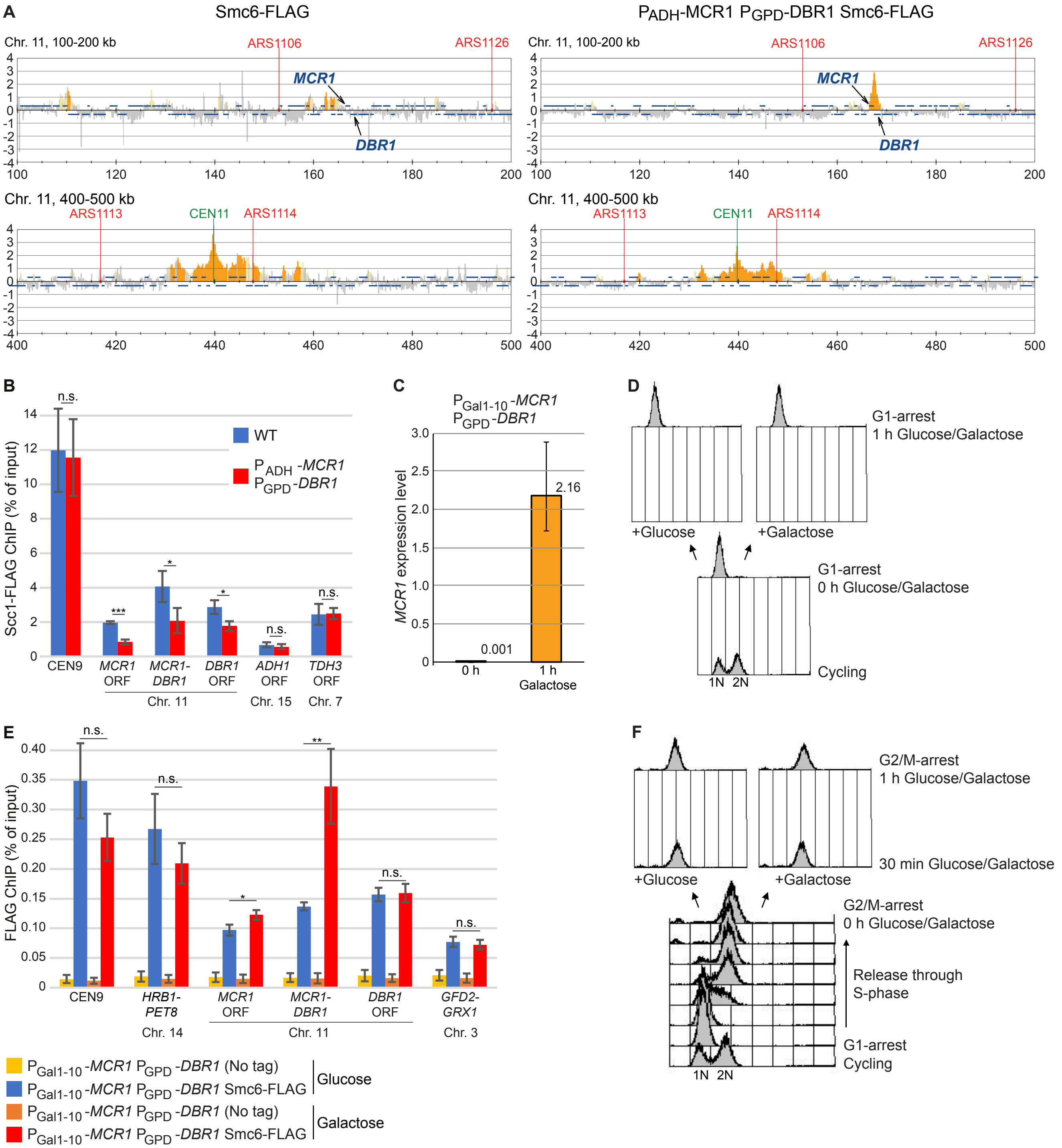
A
Smc6-FLAG
PADH-MCR1 PGPD-DBR1 Smc6-FLAG
Chr. 11, 100-200 kb
ARS1106
ARS1126
MCR1
DBR1
Chr. 11, 400-500 kb
ARS1113
CEN11
ARS1114
B
WT
PADH-MCR1 PGPD-DBR1
Scc1-FLAG ChIP (% of input)
n.s.
***
*
CEN9
MCR1 ORF
MCR1-DBR1
DBR1 ORF
ADH1 ORF
TDH3 ORF
Chr. 11
Chr. 15
Chr. 7
C
PGal1-10-MCR1
PGPD-DBR1
MCR1 expression level
2.16
0.001
0 h
1 h
Galactose
D
G1-arrest
1 h Glucose/Galactose
+Glucose
+Galactose
G1-arrest
0 h Glucose/Galactose
Cycling
1N 2N
E
FLAG ChIP (% of input)
n.s.
**
*
CEN9
HRB1-PET8
MCR1 ORF
MCR1-DBR1
DBR1 ORF
GFD2-GRX1
Chr. 14
Chr. 11
Chr. 3
PGal1-10-MCR1 PGPD-DBR1 (No tag)
PGal1-10-MCR1 PGPD-DBR1 Smc6-FLAG
Glucose
PGal1-10-MCR1 PGPD-DBR1 (No tag)
PGal1-10-MCR1 PGPD-DBR1 Smc6-FLAG
Galactose
F
G2/M-arrest
1 h Glucose/Galactose
30 min Glucose/Galactose
+Glucose
+Galactose
G2/M-arrest
0 h Glucose/Galactose
Release through S-phase
G1-arrest
Cycling
1N 2N

**Figure S6**

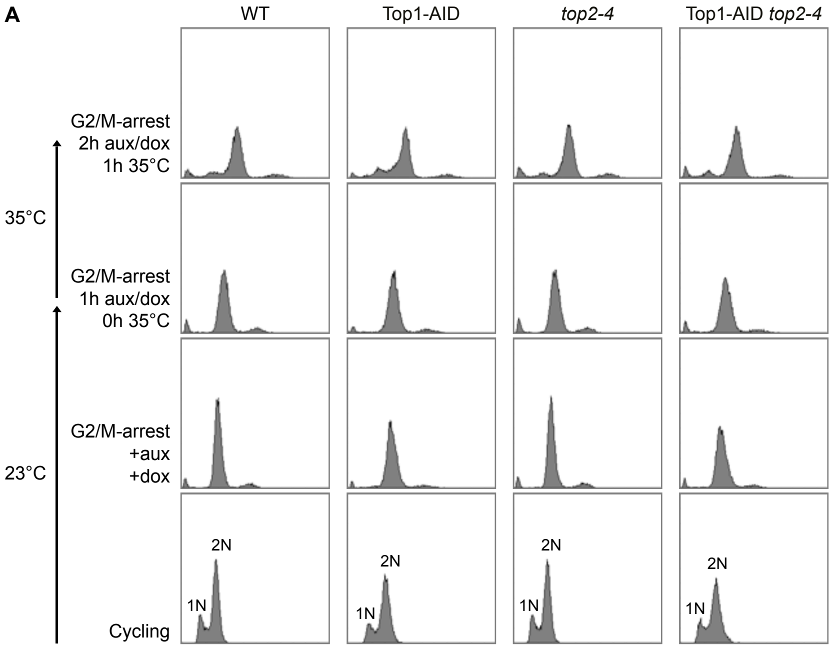

A
WT
Top1-AID
*top2-4*
Top1-AID *top2-4*
G2/M-arrest
2h aux/dox
1h 35°C
35°C
G2/M-arrest
1h aux/dox
0h 35°C
G2/M-arrest
+aux
+dox
23°C
Cycling
1N
2N

**Figure S7**

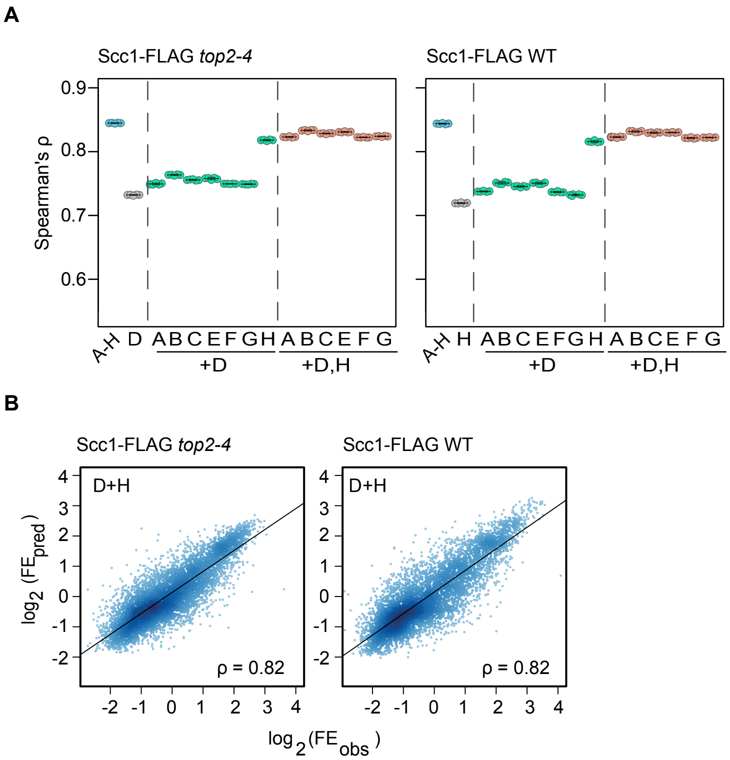
A
Scc1-FLAG top2-4
Scc1-FLAG WT
Spearman's ρ
0.9
0.8
0.7
0.6
A-H D A B C E F G H A B C E F G
+D
+D,H
A-H H A B C E F G H A B C E F G
B
Scc1-FLAG top2-4
Scc1-FLAG WT
D+H
ρ = 0.82
log2 (FEpred)
log2 (FEobs)

**Figure S8**

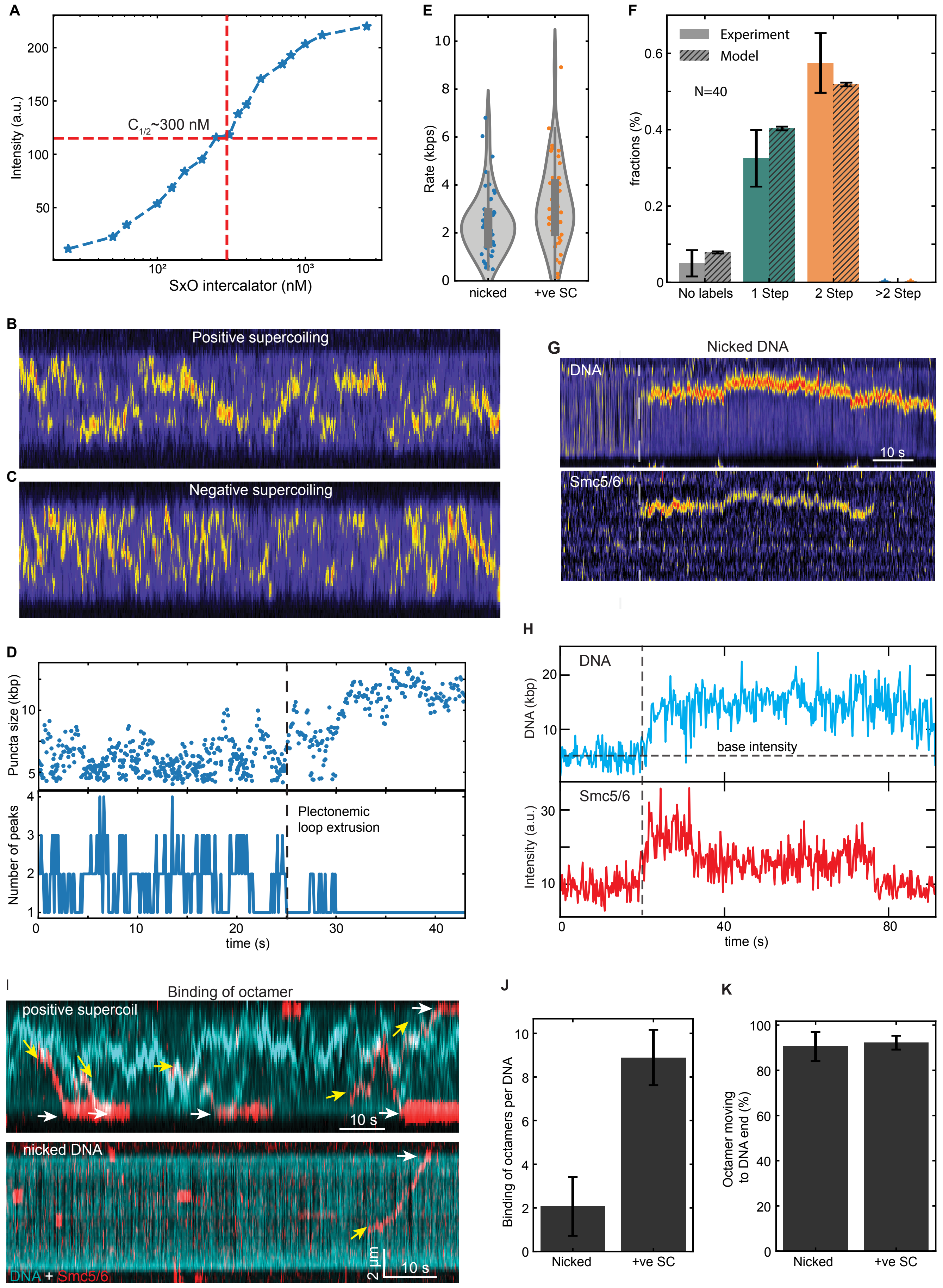

A
Intensity (a.u.)
C1/2~300 nM
SxO intercalator (nM)
E
Rate (kbps)
nicked
+ve SC
F
Experiment
Model
N=40
fractions (%)
No labels
1 Step
2 Step
>2 Step
B
Positive supercoiling
C
Negative supercoiling
G
Nicked DNA
DNA
Smc5/6
10 s
D
Puncta size (kbp)
Number of peaks
Plectonemic
loop extrusion
time (s)
H
DNA
DNA (kbp)
base intensity
Smc5/6
Intensity (a.u.)
time (s)
I
Binding of octamer
positive supercoil
10 s
nicked DNA
DNA + Smc5/6
2 µm
10 s
J
Binding of octamers per DNA
Nicked
+ve SC
K
Octamer moving
to DNA end (%)
Nicked
+ve SC

Figure S9

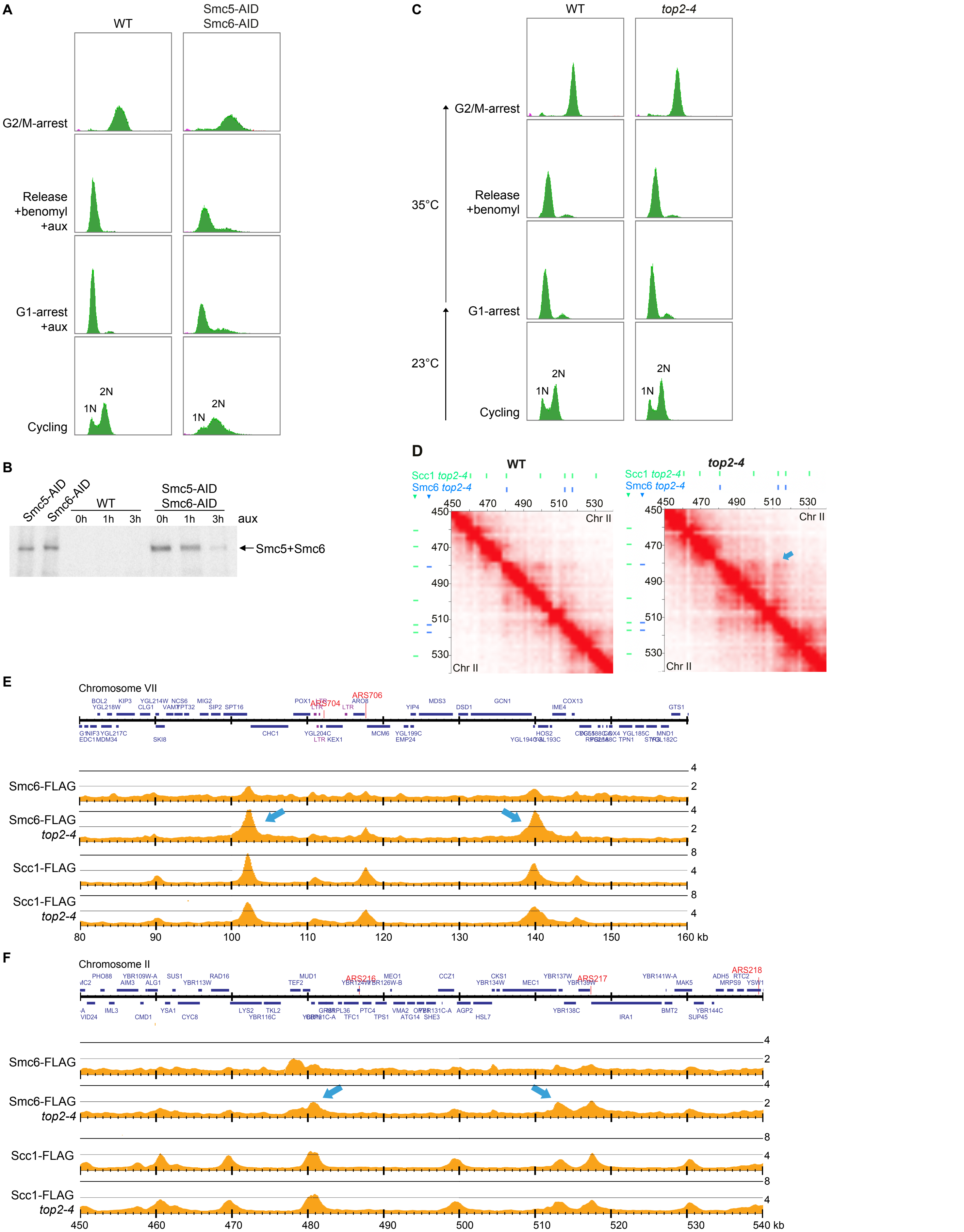
A
WT
Smc5-AID
Smc6-AID
G2/M-arrest
Release
+benomyl
+aux
G1-arrest
+aux
Cycling
1N
2N
B
Smc5-AID
Smc6-AID
WT
0h 1h 3h
0h 1h 3h
aux
Smc5+Smc6
C
WT
top2-4
G2/M-arrest
35°C
Release
+benomyl
G1-arrest
23°C
Cycling
D
Scc1 top2-4
Smc6 top2-4
450 470 490 510 530
Chr II
E
Chromosome VII
ARS704
ARS706
Smc6-FLAG
Smc6-FLAG top2-4
Scc1-FLAG
Scc1-FLAG top2-4
80 90 100 110 120 130 140 150 160 kb
F
Chromosome II
ARS216
ARS217
ARS218
Smc6-FLAG
Smc6-FLAG top2-4
Scc1-FLAG
Scc1-FLAG top2-4
450 460 470 480 490 500 510 520 530 540 kb

Figure S10

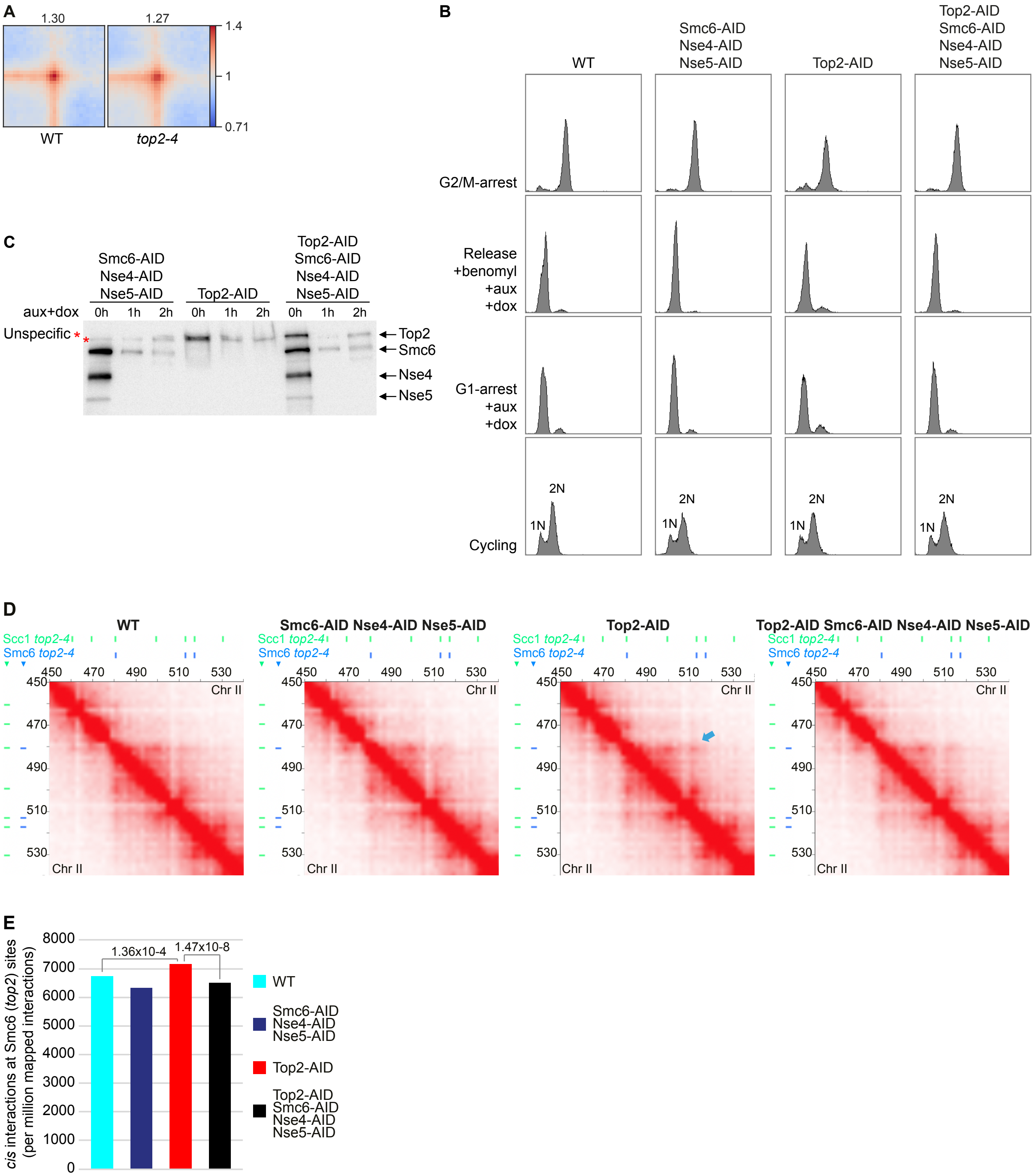
A
1.30
1.27
1.4
1
0.71
WT
top2-4
B
WT
Smc6-AID
Nse4-AID
Nse5-AID
Top2-AID
Top2-AID
Smc6-AID
Nse4-AID
Nse5-AID
G2/M-arrest
Release
+benomyl
+aux
+dox
G1-arrest
+aux
+dox
Cycling
2N
1N
C
Smc6-AID
Nse4-AID
Nse5-AID
Top2-AID
Top2-AID
Smc6-AID
Nse4-AID
Nse5-AID
aux+dox
0h 1h 2h 0h 1h 2h 0h 1h 2h
Unspecific *
Top2
Smc6
Nse4
Nse5
D
WT
Smc6-AID Nse4-AID Nse5-AID
Top2-AID
Top2-AID Smc6-AID Nse4-AID Nse5-AID
Scc1 top2-4
Smc6 top2-4
450 470 490 510 530
Chr II
E
cis interactions at Smc6 (top2) sites
(per million mapped interactions)
8000
7000
6000
5000
4000
3000
2000
1000
0
1.36x10-4
1.47x10-8
WT
Smc6-AID
Nse4-AID
Nse5-AID
Top2-AID
Top2-AID
Smc6-AID
Nse4-AID
Nse5-AID

**Figure S11**

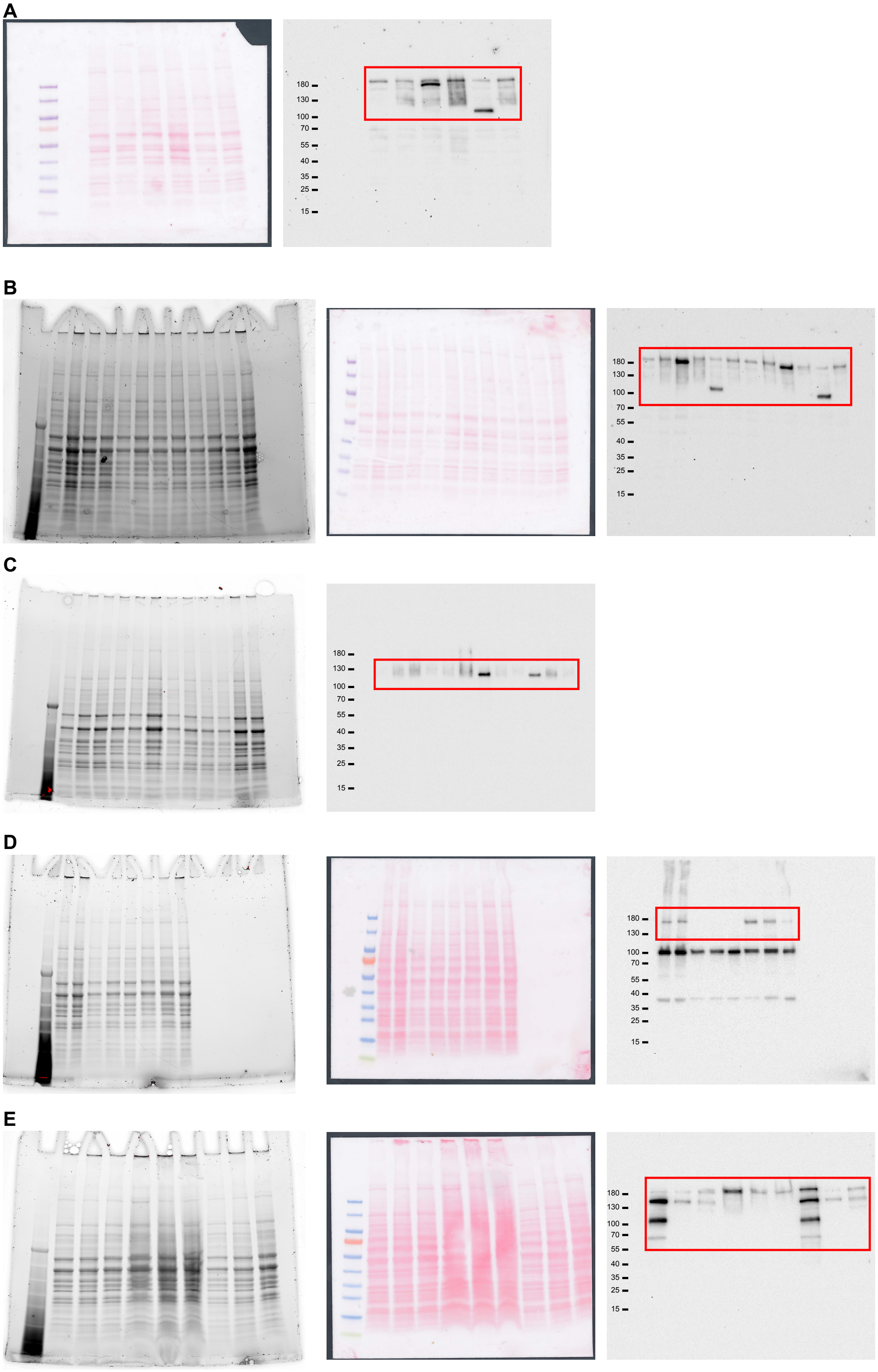

#### SUPPLEMENTAL MOVIE LEGENDS

**Supplemental Movie 1. Loop extrusion on positively supercoiled DNA by Smc5/6. Related to Figure 6.**

Kymographs show how dynamic positive plectonemic supercoils, observed as transient local maxima, are gathered into a single intensifying spot by hexameric Smc5/6.

**Supplemental Movie 2. Loop extrusion on positively supercoiled DNA by fluorescently labelled Smc5/6. Related to Figure 6.**

Kymographs show that Nse4-labelled hexameric Smc5/6 (middle) binds to the dynamic positively supercoiled plectomemes and gather them into a single intensifying spot by loop extrusion (top). Overlay (bottom).

**Supplemental Movie 3. Smc5/6 loop extrudes from the tip to the base of a positively supercoiled plectoneme. Related to Figure 6.**

Kymographs show that under buffer side-flow conditions, Smc5/6 (middle) binds to the top of the positively supercoiled DNA plectoneme (top), and loop extrudes to the loop base. Overlay (bottom).
